# Estimating the effective population size across space and time in the Critically Endangered western chimpanzee in Guinea-Bissau: challenges and implications for conservation management

**DOI:** 10.1101/2024.10.07.616982

**Authors:** Maria Joana Ferreira da Silva, Filipa Borges, Federica Gerini, Rui M. Sá, Francisco Silva, Tiago Maié, Germán Hernández-Alonso, Jazmín Ramos-Madrigal, Shyam Gopalakrishnan, Isa Aleixo-Pais, Mohamed Djaló, Nelson Fernandes, Idrissa Camará, Aissa Regalla, Catarina Casanova, Mafalda Costa, Ivo Colmonero-Costeira, Carlos Rodríguez Fernandes, Lounès Chikhi, Tânia Minhós, Michael W. Bruford

## Abstract

Effective population size (*N_e_*) is a key concept in evolutionary and conservation biology. The western chimpanzee (*Pan troglodytes verus*) is a Critically Endangered taxon. In Guinea-Bissau, chimpanzees are mainly threatened by habitat loss, hunting and diseases. Guinea-Bissau is considered a key area for its conservation. Genetic tools have not yet been applied to inform management and no estimates of *N_e_* have been obtained. In this study, we use country’s range-wide microsatellite data and five whole-genome sequences to estimate several *N_e_* and infer the recent and ancient demographic history of populations using different methods. We also aim to integrate the different *N_e_* estimates to improve our understanding of the evolutionary history and current demography of this great ape and to discuss strengths and limitations of each estimator and their complementarity in informing conservation decisions. Results from the PSMC method suggest a large ancestral *N_e_*, likely due to ancient structure over the whole subspecies distribution until approximately 10-15,000 years ago. After that, a change in connectivity, a real decrease in size or a combination of both occurred, which reduced the then still large ancestral population to a smaller size (MSVAR: ∼10,000 decreasing to 1,000-6,000 individuals), possibly indicating a fragmentation into coastal and inner subpopulations. In the most recent past, contemporary *N_e_* is below or close to 500 (GONE: 116-580, NeEstimator: 107-549), suggesting a high risk of extinction. The populations at coastal Parks may have been small or isolated for several generations whereas the Boé Park one exhibit higher long-term *N_e_* estimates and can be considered a stronghold of chimpanzee conservation. Through combining different types of molecular markers and analytical methodologies, we try to overcome the limitations of obtaining high quality DNA sampling from wild threatened populations and estimate *N_e_* at different temporal and spatial scales, which is crucial information to make informed conservation decisions at local and regional scales.

## 1. Introduction

The concept of effective population size (*N_e_*) is central in evolutionary and conservation biology and has important practical applications in conservation management (Frankham et al., 2010, Hoban et al., 2022, Waples, 2022). *N_e_* is considered as probably the most important metric to understand and predict both the populations’ short-term risk of extinction by inbreeding depression and their long-term potential to adapt to environmental changes (Hoban et al., 2020, Hoban et al., 2022). *N_e_* is also one of the best-studied metrics for applying minimal viable population thresholds and identifying populations of conservation concern (Frankham, 2005, Jamieson & Allendorf, 2012, Frankham et al., 2014). Moreover, *N_e_* has many practical applications in wildlife management and conservation planning, such as designing human-induced translocations and relocation of populations (Luikart et al., 2010, O’Brien et al., 2022, Waples, 2022, Waples 2024). For instance, *N_e_* has been considered one of the four genetic *Essential Biodiversity Variables* (EBVs, summary measures of biodiversity), which are designed to monitor changes in biodiversity over time and space (Hoban et al., 2022).

*N_e_* seems easy to understand and to compute using genetic data (Allendorf et al., 2010). However, it is perhaps one of the most difficult and error-inducing concepts to grasp in population genetics. One reason for this is that *N_e_* is a single number that aims at summarizing a usually highly complex situation, whether we are interested in the demographic history or on the recent dynamics of a species (Chikhi et al., 2010, 2018, Wakeley, 1999, Waples 2022). Furthermore, the estimation and practical integration of the *N_e_* parameter into conservation management and policies is advancing slowly even in regions with high biodiversity and iconic and endangered species (Bertola et al., 2024), such as great apes. This is related to a number of factors concerning feasibility and low financial resources for population genetic studies to estimate *N_e_* and a low reliability of results when the estimation is carried out for non-model species not meeting the assumptions of population genetic models (e.g., Bertola et al., 2024; Waples 2022).

The concept of *N_e_* was introduced by Sewall Wright in 1931 (Wright, 1931). *N_e_* quantifies the rate of genetic change (e.g., drift of allele frequencies) of real populations in reference to the Wright-Fisher (WF) idealized population (Wang et al., 2016). WF populations are assumed to have equal sex ratio, constant size and non-overlapping generations, no sexual or natural selection and in which genetic drift is considered the only evolutionary force changing gene frequencies across generations together with mutations (Conner & Hartl, 2004). *N_e_* is thus the size of an idealized WF population with the same properties of genetic drift as the real (more complex) population under consideration (Wang et al., 2016).

While the concept seemed straightforward, it was later realized that one could identify different types of *N_e_* depending on the property of interest that *N_e_* was supposed to summarize (*e.g*., Ryman et al., 2019). For instance, one can define the *N_e_* generating the same rate of inbreeding as the real population of interest (*i.e.*, the probability that a pair of homologous genes in an individual came from the same parent in the previous generation, which was denoted as the inbreeding effective size, *N_eI_*) or the *N_e_* generating the same rate of change in variance of gene frequencies (denoted as the variance effective size, *N_eV_*) (Wang et al., 2016). More recently, the concept of coalescent *N_e_* was also defined to identify the *N_e_* that can explain the patterns of diversity observed in present-day populations under simple demographic models, typically assuming panmixia over long periods of time. This concept has itself been extended by allowing *N_e_* to change through time. Note that under the latter case, there is not one *N_e_* but rather a succession of *N_e_* values, which may thus lead to apparent contradictions between methods estimating one *N_e_* and those estimating a succession of *N_e_* values. Under a standard constant-size WF model, all the different *N_e_* concepts are expected to be the same. However, this is not necessarily the case in real-world situations, where populations are rarely panmictic or at mutation-drift equilibrium. Real populations have likely gone through complex demographic histories involving expansions and contractions, related to environmental changes and fragmentation of habitats (Wang et al., 2016; Ryman et al., 2019). In addition, theoretical work suggests that there may be demographic models for which some *N_e_* cannot be defined (Sjödin et al., 2005). The point we wish to make here is that depending on the research questions asked, one may obtain very different answers. The fact that we obtain different values should be seen as an indication that the species of interest may not be easily summarized by a single *N_e_* number, and that the different estimates obtained might all be useful for devising conservation strategies that account for both the ongoing dynamics of the species but also for its demographic history.

The western chimpanzee (*Pan troglodytes verus*, Schwarz, 1934) is one of the four currently recognized subspecies of chimpanzees *P. troglodytes*. Its range extends from Senegal in the west to Ghana in the east (Fig. 1a). The subspecies presently occurs in the following eight West African countries: Côte d’Ivoire, Ghana, Guinea, Guinea-Bissau, Liberia, Mali, Senegal and Sierra Leone, and has most likely disappeared from Benin, Burkina Faso, and Togo (Campbell & Houngbedji, 2015, Ginn et al., 2013, IUCN SSC Primate Specialist Group 2020). This subspecies has been classified as Critically Endangered by the International Union for Conservation of Nature (IUCN) (Humle et al., 2016). The population of *P. t. verus* is estimated to have decreased its abundance by 80% between 1990 and 2014 and to have reached a global size between 15,000 and 65,000 individuals (Kühl et al., 2017) or 52,811 individuals (95% CI 17,577–96,564) as more recently estimated by Heinicke et al. (2019a). The subspecies conservation status is expected to deteriorate in the next decades considering that the great majority of the western chimpanzees currently live outside protected areas and within 5 km of an infrastructure (Heinicke et al., 2019a), and it is predicted high rates of deforestation until 2050 for its West African range (Palminteri et al., 2018). Moreover, the western chimpanzee is threatened by hunting to supply the trade of wild meat, live animals and body-parts, and by diseases (Humle et al., 2016; IUCN SSC Primate Specialist Group 2020, Sá et al., 2012). The western chimpanzee has low genetic diversity when compared to the other *P. troglodytes* subspecies, and two recent studies have suggested that *N_e_* could be in the order of 17,378 breeding individuals (de Manuel et al., 2016; Fontsere et al., 2022), even if this number should be interpreted with care.

**Figure 1.**
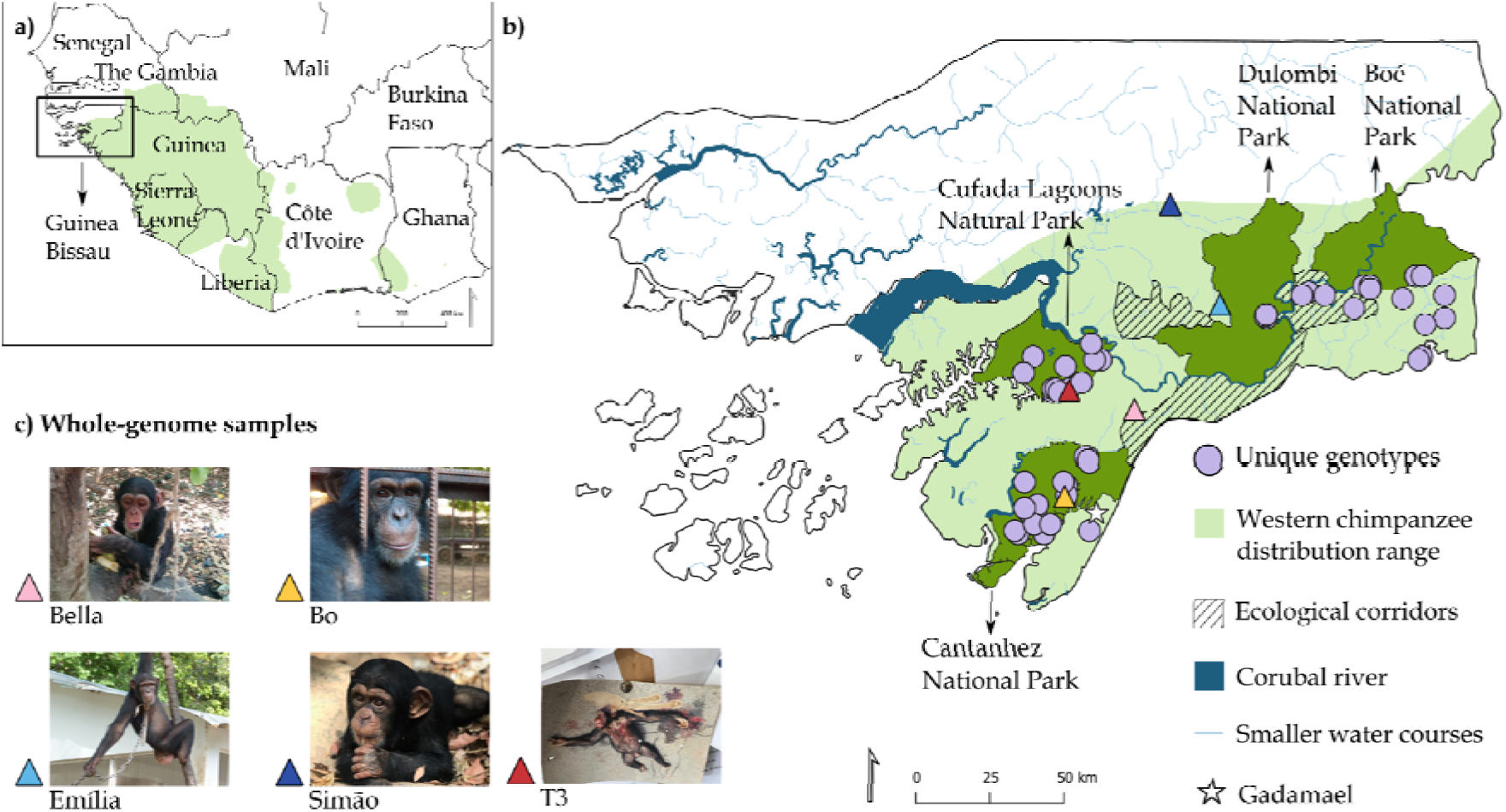
a) Distribution of the western chimpanzee (*Pan troglodytes verus*) in West Africa and b) the location of the study area in Guinea-Bissau. Unique genotypes for 10 microsatellite loci (represented by purple circles, N=143) were obtained from fecal samples collected non-invasively in four parks located in southern Guinea-Bissau and encompassing the national range of chimpanzees – Cufada Lagoons Natural Park (CLNP), Cantanhez National Park (CNP), Dulombi National Park (DNP) and Boé National Park (BNP). The location of ecological corridors is also indicated. Gadamael area (referred to in table 1) is also mapped and represented as a white star. c) Pictures show the four confiscated chimpanzees and one road-killed individual that were sampled to generate whole-genome sequencing data. Blood samples were drawn from the confiscated individuals during placement in a sanctuary abroad. One tissue sample was obtained from one road killed minutes after fatality. The location of confiscated individuals in the map in b) (represented by triangles) is estimated and reflects information on the individual’s origin obtained from national authorities (*e.g.,* Bo from CNP) or where individuals were found in private premises. Photo credits by MJFS, H. Foito (European Union Embassy Bissau), C. Casanova, L. Almeida and I. Camará.

Guinea-Bissau (GB) (area: 36,125 km, population: 2,08 million) is an important biodiversity hotspot holding populations of emblematic and threatened species, such as leopard (*Panthera pardus*, Linnaeus, 1758), lion (*Panthera leo*, Linnaeus, 1758), elephant (*Loxodonta cyclotis*, Matschie, 1900), saltwater hippopotamus (*Hippopotamus amphibius*, Linnaeus, 1758), manatee (*Trichechus manatus*, Lineu, 1758) (Brugiere et al., 2005; Palma et al., 2023), and ten confirmed primate species, including the western chimpanzee, colobus monkeys (*Piliocolobus temminckii*, Kuhl, 1820 and *Colobus polykomos*, Zimmermann, 1780) and the sooty mangabey (*Cercocebus atys*, Audebert, 1797) (Bersacola et al., 2018; Ferreira da Silva et al., 2020, Minhós et al., 2023). Biodiversity conservation has been considered a driver for economic development and consequently, a great effort has been made by national agencies to formally protect a network of areas, which includes six Parks and three ecological corridors in mainland Guinea-Bissau and the Bijagós archipelago (Fig. 1), covering almost 26% of the country size (https://ibapgbissau.org/areas-protegidas/). Nevertheless, national conservation management needs some important improvements, such as a corrected list of the species occurring in the country, its present range (*e.g.*, Ferreira da Silva et al., 2020), and baseline estimates of demographic parameters of threatened species, such as population size, to inform prioritization of areas to conserve.

The western chimpanzee (*Dari,* in Guinea-Bissau Creole) occurs in the southern part of the country, mainly south of the Corubal River (Bersacola et al., 2018; Carvalho et al., 2013) (Fig. 1b). Chimpanzees were erroneously declared extinct and were rediscovered in the 1990s (Gippoliti & Dell’Omo 1996, 2003). In GB specifically, the main conservation threats faced by the subspecies are habitat loss and fragmentation, hunting to supply the trade of live individuals and retaliatory killing during crop-raiding (Hockings & Sousa, 2013). Please note that the trade of body parts, such as skins and bones, for traditional medicine practices is observed in the capital city markets, although the national origin of these specimens has not been confirmed (Sá et al., 2012), and the trade and consumption of chimpanzee meat does not seem to occur (Ferreira da Silva et al., 2021, Minhós et al., 2013, van Laar 2010), contrary to what happens in other countries (*e.g*., Côte d’Ivoire; Caspary et al., 2001). Chimpanzee meat in GB is considered non-edible by locals because this primate species is considered to have high resemblance with humans (Amador et al., 2014, Gippoliti & Dell’Omo 2003, Karibuhoye, 2004, Sousa et al., 2017). In contrast, trade of live infant chimpanzees is a very frequent phenomenon (Ferreira da Silva & Regalla, under review, Hockings & Sousa 2013) and may be a significant threat since its capture involves killing the adults (Ferreira da Silva et al., 2021). In addition, chimpanzees appear to be threatened by the propagation of diseases such as leprosy (*Mycobacterium leprae*), which has been detected in several communities of Cantanhez National Park (CNP, Fig. 1) (Hocking et al., 2021). Past studies reporting a high prevalence of parasites shared with humans suggest that habitat disturbance plays a role in the transmission and persistence of pathogens (Sá et al., 2013). GB is an important area in West Africa for the conservation of *P. t. verus*. Specifically, i) the coastal areas of GB together with the ones in Republic of Guinea are considered a priority region (Kormos & Boesch, 2003; IUCN SSC Primate Specialist Group 2020), ii) the protected areas of CNP, Dulombi National Park (DNP) and Boé National Park (BNP) (Fig. 1) are considered areas of high value for conservation (Heinicke et al., 2019a), and iii) the Boé region, in the inner part of the country, is one of eight sites across the subspecies distribution that is classified as exceptionally stable or of high-density (Heinicke et al., 2019b).

Historically, the overall population in the country has been suggested to be between 600 and 1,000 (Gippoliti & Dell’Omo, 2003) and more recently estimated as 1,908 individuals (95% confidence interval: 923–6,121 individuals Heinicke et al., 2019a). Improved representative surveys have been recommended for GB given the large confidence intervals of estimates (Heinicke et al., 2019a). The size of local populations has been evaluated for most of the protected areas where chimpanzees occur using various indirect methods (Table 1). Although the estimates from the different studies cannot be compared directly, Cufada Lagoons Natural Park (CLNP) is highlighted as the population with the lowest density, whereas the Boé region stands out as the one displaying the topmost density (reaching > 6 individuals per km^2^, Table 1). Currently, there is no size or density assessment for DNP or for other populations outside areas with formal protection, such as ecological corridors (but see the exception of Gandamael, Table 1, Fig. 1b), although it is estimated that approximately 35% of chimpanzees live outside a park in GB (Heinicke et al., 2019a). Genetic tools have not yet been applied to inform the conservation of the western chimpanzees in GB. Little is known about the genetic diversity or the amount of genetic isolation between populations (but see Borges, 2017 and Gerini, 2018) and no estimates of *N_e_* have been obtained to date.

**Table 1.** Compilation of results from past studies estimating the density of local populations of chimpanzees in Guinea-Bissau.

| Site | Reference | Method | Estimated density (ind./km<sup>2</sup>) | Estimated number of individuals |
| --- | --- | --- | --- | --- |
| Cantanhez National Park | Sousa J. 2009 | nest count method for density estimation | 1.937 and 2.340 | *2,070 and 2,454 |
|  | Torres et al., 2010 | modeling | MI | 376 and 2632 |
|  | Hockings et al., 2013 | MI | 3.00 | MI |
| Cufada Lagoons Natural Park | Carvalho et al., 2013 | nest count method for density estimation | 0.22 | 137 (95% C.I. 51–390) |
|  | Sousa et al., 2013 | marked nest count method | 0.79 (CI 95 %: 0.61–1.04) | 300 (CI 95 %: 230–390) |
| Gadamael | Sousa F. (2009) | nest count method for density estimation | 0.897 | 33 |
| Boé National Park | Schwarz et al., 2007 | Interviews | MI | 710 |
|  | Dias et al., 2018++ | nest count method for density estimation | 0.38 | 18 |
|  | Wenceslau, 2014 | nest count method for density estimation | 1.8 | MI |
|  | Binczik et al., 2017 | standing crop nest count method | 0.77 (95% CI 0.45–1.34) | 1,465 to 4,415 |
|  | Heinicke et al., 2019a | modeling | 6.76 | MI |
| Overall population | Heinicke et al., 2019a | modeling |  | 1,908 (923–6121 95% CI) |

In this study, we use geographically broad genetic data and genomic data from multiple wild-born individuals to estimate the *N_e_* of the western chimpanzee population in GB. We aim to i) estimate *N_e_* and infer the recent and ancient demographic history of populations using different methods as applied to microsatellite loci and whole-genome sequence data (WGS), ii) integrate the different estimates to improve our understanding of both the evolutionary history and current demography of the western chimpanzees, iii) discuss the strengths and limitations of each *N_e_* estimator method and their complementarity in informing conservation decisions for long-lived organisms, and iv) discuss the implications of the results for the conservation management of this emblematic species in GB.

## 2. Materials and methods

### 2.1 Study area

The study area covers a large proportion of the chimpanzee range in GB (Gippoliti & Dell’Omo 2003) (Fig. 1b), encompassing an area of approximately 6,000 km. Sampling of biological material was carried out in four geographically-distinct and formally protected areas – 1. Cantanhez National Park (CNP, 1,067.67 km^2^), 2. Cufada Lagoons Natural Park (CLNP, 890 km^2^), 3. Dulombi Natural Park (DNP, 1,600.96 km²) and 4. Boé National Park (BNP, 1,552.95 km^2^) (https://ibapgbissau.org/areas-protegidas/). Chimpanzees were known to be present at these areas prior to our study (Gippoliti & Dell’Omo 2003; Bersacola et al., 2018).

### 2.2 Microsatellite loci dataset

We generated a dataset of 143 unique genotypes for 10 microsatellite loci derived from non-invasive fecal samples (Borges, 2017, Gerini, 2018) (Fig. 1b). Eighty-five genotypes correspond to samples collected between 2015 and 2017 in CLNP (N=38), BNP (N=34), and DNP (N=13) and the remaining consisted of previously determined genotypes from CNP (N=58) (Sá, 2013) (Fig. 1b). Fecal samples were collected fresh and from unhabituated and unidentified individuals, in sites used by chimpanzee groups for sleeping, foraging and drinking. The techniques and methods to preserve the fecal samples until DNA extraction are described in Ferreira da Silva et al. (2014). DNA extraction was carried out using two methods: i) the QIAamp®DNA Stool Mini Kit (QIAGEN®) at MWB research group laboratory facilities at School of Biosciences, Cardiff University, UK (Sá, 2013) and ii) the CTAB method (Vallet et al., 2008, adapted by Quéméré et al. 2010) for samples collected between 2015-2017, which were extracted at *Instituto Gulbenkian de Ciência* (IGC, Oeiras, Portugal) laboratory facilities. The procedures to avoid contamination by exogenous DNA are described elsewhere (Ferreira da Silva et al., 2014). DNA samples were identified to the species level using a mitochondrial DNA hypervariable region I fragment (approximately 600 base pairs, using primers L15926 and H16555, as described in Sá, 2013). Consensus sequences were derived from forward and reverse sequencing by visual comparison using Geneious Pro v.4.8.5 (Biomatters, Biomatters Ltd, New Zealand). Standard Nucleotide BLAST in NCBI (http://www.ncbi.nlm.nih.gov/) was used to identify accessions closely related to the generated sequences and confirm that samples were from *P. troglodytes verus* (*i.e*., GenBank Accession code D38113). Allele size standardization between datasets was carried out using re-extraction and re-analyses of DNA extracts of five samples included in Sá (2013) together with the novel samples analyzed in Borges (2017) and Gerini (2018). Allele scoring followed previously described procedures to guarantee minimal impact of allelic dropout and false alleles errors: four replicates were carried out per sample and the rules to reach a consensus genotype were determined per locus (Ferreira da Silva et al., 2014). The consensus genotype was classified according to the Quality Index (QI, Miquel et al., 2006), and genotypes with a mean across loci below 0.55 were excluded from the dataset. The probability of identity (PI) and the probability of identity between siblings (PIsibs) (Waits et al., 2001), estimated using GenAIEx v.6.503 (Peakall & Smouse, 2006), was of 1.5 × 10^−11^ and 8.9 × 10^−05^, respectively, which in principle allows to distinguish between unique genotypes using six loci. We could not find genotyping errors (typing errors, large-allele dropout, and locus-specific deficiency in heterozygotes due to null alleles) using MicroChecker v.2.2.3 (van Oosterhout et al., 2006) apart from locus D2S1326 for CLNP, which showed excess of homozygotes. We retained the locus in the final dataset as we found no significant departures from Hardy-Weinberg equilibrium per locus using the Bonferroni correction when geographic populations were analyzed separately. The population of chimpanzees in GB does not display significant population structure when assessed using individual-based Bayesian algorithms (*e.g.,* STRUCTURE) (Borges, 2017 estimated K=1).

### 2.3 Genomic data

#### 2.3.1 Sampling

Whole-genome sequences were produced from biological material collected from wild born chimpanzees: one road-killed (tissue sample T3-Chimp collected in 2011) and four individuals (blood samples from Bo, Bella, Simão and Emilia chimpanzees, collected between 2018 and 2019) confiscated by the Institute for Biodiversity and Protected Areas (IBAP) from private premises. We obtained the information that the individuals were caught by hunters in different sites within GB (*e.g*., Bo was originally from CNP, Bella was confiscated in Quebo but probably originated from CNP, Emilia was from DNP, and Simão was living in Bafatá but traded in Quebo, Fig. 1). Blood was collected as part of the placement of the individuals in a sanctuary abroad (Sweetwaters Chimpanzee Sanctuary, Ol Pejeta, Kenya, Ferreira da Silva & Regalla, in review^3^). The blood samples were drawn by a wildlife veterinarian (P. Melo, vet_natura, https://www.vetnatura.pt/) for health screening and as part of a parasites and virus detection procedure prior to translocation (Melo et al., 2018). Samples were collected in 5 mL collection tubes filled up with the anticoagulant ethylenediamine tetraacetic acid (EDTA) and preserved fresh until DNA extraction. The road-killed individual was found in the road next to CLNP (Fig. 1b) and a sample of muscle tissue was collected and preserved in 98% ethanol up to DNA extraction.

#### 2.3.2 DNA extraction and data production

DNA was extracted from the five samples adapting the method by Vallet et al., (2008). We used 500 µL of each blood sample and about 10 mg of tissue from the road-killed individual. The details of this two-day DNA extraction protocol can be found in Supplementary material S1. We tested the quality of DNA extractions in 2% agarose gels and quantified DNA concentration using a Nanodrop microvolume spectrophotometer (ThermoFisher Scientific) (Supplementary material S2). Laboratory procedures took place at the IGC, and extractions were carried out in a biological safety cabinet in a Biosafety Level 2 dedicated room. Library preparation and sequencing were performed by Macrogen at a coverage of 30-15x using the Illumina Hiseq X and TruSeq platforms.

#### 2.3.3 Whole-genome sequence (WGS) data assembly, Mapping and Genotype Calling

After all samples passed quality control tests, we used the BAM pipeline from PALEOMIX to process the sequences for downstream analysis at the Globe Institute’s (University of Copenhagen, Demark) High-Performance Computing (HPC) cluster. This pipeline trims adapter sequences, filters low quality reads, removes PCR duplicates, and aligns reads and maps them to a reference genome (Schubert et al., 2014). We used a “makefile” (.yaml file), that allows the specification of the tasks to be performed, BWA as the aligner software and the algorithm “mem”. The BWA-mem algorithm shows great performance with sequencing errors and is most adequate for short reads, as it is the case of this study (Li, 2013). The “MinQuality” parameter was used to exclude reads with a mapping quality (or Phred score) below zero (Schubert et al., 2014).

For the genotype calling, we first selected variants with a minimum Phred quality score of 20, using the HaplotypeCaller algorithm in GATK (version 4.2.0.0; Poplin et al., 2017). HaplotypeCaller uses the input data to calculate the likelihood of each genotype per sample, and then assigns the most likely genotype to that sample. Through the application of the ‘SelectVariants’, only sites with SNPs were selected. Lastly, *vcftools* (Danecek et al., 2011) was used to remove indels, namely the sites with a missing proportion higher than 0.9, a Phred quality score equal to or lower than 30, and genotypes with depth values lower than 5 and higher than 100. A table describing the WGS data summary statistics for each sample, such as coverage, observed number of homozygous genotypes, expected number of homozygous genotypes and inbreeding coefficient (F), can be found in Supplementary material S3.

As a quick assessment of the possible presence of genetic structure among the sampled individuals, we performed an analysis with the STRUCTURE software version 2.3.4 (Pritchard et al., 2000) using the admixture model and assuming correlated allele frequencies. The parameter set consisted of a burnin period of 50,000 steps, followed by 200,000 iterations, and 10 runs for each number of clusters (Van Wyngaarden et al., 2017). The results indicated evidence for a single panmictic cluster (K=1) as the best clustering solution to explain the observed genetic variation across individuals, *i.e.* absence of genetic structure (Silva, 2024).

### 2.4 Effective population size estimation and demographic history

#### 2.4.1 The PSMC (pairwise sequentially Markovian coalescent) and the IICR: principles of demographic inference

The PSMC method of Li and Durbin (2011) was applied to the nuclear genomes of the five individuals for which tissue and blood samples were obtained. The PSMC uses the information from the distribution of heterozygous sites along the genome of a single diploid individual (or two haploid genomes) and produces a curve where the x-axis represents time usually represented in a log-scale, and the y-axis is often interpreted as representing the effective population size. Proper scaling of the PSMC in years requires the use of estimates of generation time, mutation, and recombination rates.

PSMC version 0.6.5-r67 (Li & Durbin, 2011) (available at http://github.com/lh3/psmc) was run on each individual genome using the following settings: -N25 -t15 -r5 -p “4+25*2+4+6”. Individual consensus sequences were generated using the *mpileup*, *bcftools* and *vcfutils.pl* (*vcf2fq*) pipeline from SAMTOOLS v. 1.16, with minimum read depth (-d) set to five and maximum read depth (-D) set to 30. The consensus sequence was converted into a fasta-like format using the *fq2psmcfa* program, provided in the PSMC package, with the quality cut off (-*q*) set to 20. We assumed a mutation rate (*μ*) of 1.2 × 10^−8^ per base pair per generation and a generation time of 25 years (Venn et al., 2014; Besenbacher et al., 2019; Chintalapati & Moorjani, 2020). To quantify the variance in PSMC curves, we performed 10 bootstraps per individual, following the re-sampling protocol suggested by the authors. The inferred demographic histories for the five analyzed individuals were plotted in a single figure using Ghostscript 9.16 and Gnuplot 5.4.0. The PSMC plots are usually interpreted in terms of *N_e_* changes but can also be interpreted in terms of connectivity changes (see Discussion).

#### 2.4.2 MSVAR analysis of microsatellite loci data

We also used the Bayesian likelihood-based approach of Storz & Beaumont (2002), as implemented in the MSVAR 1.3 software. This approach assumes a simple model of exponential population size change (allowing for either growth or decline) from an ancient population of size *N_1_* to a present-day population of size *N_0_*. In practice, the method uses a Monte Carlo Markov chain algorithm to estimate the posterior probability distribution of *N_0_* and *N_1_* and of the time at which the population started to increase or decrease (*T*, in years, assuming that a generation time is given), and the *per* locus mutation rate (μ). We conducted four independent runs with different initial values and varying sets of priors and hyperpriors to reflect assumptions of either constant population size (*N_0_* = *N_1_*), population decline (*N_0_* < *N_1_*), or population growth (*N_0_* > *N_1_*) and therefore control for the impact of the priors on the posterior distributions (Supplementary material, Table S4).

Analyses were run using a series of datasets, to discard the possibility that the presence of related individuals, the sampling scheme and genetic structure could impact the inferred demographic histories, and assuming a generational span of 25 years (following Langergraber et al., 2012). We aimed to recover the demographic history of western chimpanzees by analyzing samples from i) GB as one population (N = 143 genotypes), ii) by park (CNP N = 58, CLNP N = 38, DNP N = 13 and BNP N = 34); iii) a dataset formed by unrelated individuals (N = 121, see below how relatedness was estimated); and iv) five “random datasets” obtained by randomly selecting 58 samples, which correspond to the largest dataset from a single area (*i.e.,* CNP). Datasets iii and iv were used to test the influence of the presence of highly related individuals and differences in sample sizes across datasets in demographic estimates, respectively. Analyses were also run for each geographic population since coalescent theory predicts that when populations are structured, samples obtained from one population will tend to exhibit signals of bottlenecks, whereas samples obtained across many demes will tend to have a much weaker bottleneck signal (Beaumont, 2004, Wakeley, 1999), as shown on simulated data (Chikhi et al., 2010).

To estimate relatedness between individuals, we calculated the correlation coefficient between the observed and simulated values of relatedness (100 pairs) for the Milligan (2003) and Wang (2007) likelihood estimators using the *related* R package (Pew et al., 2015). We also estimated relatedness between pairs of individuals in the overall dataset and per protected area (CNP, CLNP, DNP, and BNP) using the ML-Relate likelihood method (Kalinowski et al., 2006). We performed 100,000 simulations to identify dyads with a likely relationship of Parent-Offspring or Full-Siblings and r > 0.5 significantly different (p < 0.05) from dyads with a likely relationship of Half-Siblings and Unrelated. For dyads identified using the full dataset and the protected area dataset, one genotype of the dyad, the one which displayed lower QI, was removed from the dataset.

The individuals present in the five “random databases” were selected from the dataset i) using *runif* function in R.

Each run in MSVAR included 300,000 thinning update steps and 30,000 thinning intervals, totaling 9 × 10^9^ steps. We discarded the first 10% of each simulation to eliminate the influence of initial conditions on parameter estimation (burn-in). We verified convergence between runs both visually and using Brooks, Gelman, and Rubin Convergence Diagnostic test (Gelman & Rubin, 1992; Brooks & Gelman, 1998) conducted in R version 2.11.1 (R Development Core Team 2010) using the package BOA version 1.1.7 (Smith, 2007).

#### 2.4.3 Linkage disequilibrium-based estimation of current *N_e_* and recent demographic history from genomic data

We used GONE (Genetic Optimization for *N_e_* Estimation) (https://github.com/esrud/GONE) that implements a genetic algorithm to infer the recent demographic history of a population from single nucleotide polymorphism (SNP) data of one contemporary sample (Santiago et al., 2020). The method can infer the demographic history of a population within the past few hundreds of generations, with the authors stressing that the greatest reliability and resolution is for the last 100 generations (Santiago et al., 2020). It uses the observed spectrum of linkage disequilibrium (LD) between pairs of loci over a wide range of genetic distances (implicit recombination rates) and has been validated by simulation under different demographic scenarios and for small sample sizes (*i.e.,* n=10, Santiago et al., 2020). These simulations suggest that even when GONE is not able to accurately infer the recent *N_e_* trajectory, it can estimate the current *N_e_* relatively well. The simulations also suggested that not considering LD data between more distant loci (*e.g.*, with scaled recombination rates > 0.05) in the analyses allows for better estimates of *N_e_*, particularly when sample sizes are small or when there is population structure and migration rates between subpopulations are low (Santiago et al., 2020). Thus, the authors of the method recommend using a maximum recombination rate of 0.05, this being the default value of this parameter in GONE. Accordingly, we used this value and default settings for all other software parameters, such as no minimum allele frequency cutoff and 40 independent replicate runs, with the *N_e_* point estimate for each generation being the geometric mean of the values of the replicates. Analyses were performed using 57,650 genome-wide autosomal SNPs with genetic map information. As suggested in Santiago et al. (2020), we obtained an empirical 95% confidence interval by running GONE on 20 random replicates of 57.5K SNPs sampled from the whole-genome sequences. Replicates were generated by variant thinning in PLINK 1.9 (www.cog-genomics.org/plink/1.9/) (Chang et al., 2015).

#### 2.4.4 Linkage disequilibrium estimation of contemporary *N_e_* from microsatellite data

We used NeEstimator 2.1 (Do et al., 2014) to estimate contemporary *N_e_* from microsatellite loci data using the bias-corrected version of the method based on LD (Hill, 1981; Waples, 2006; Waples & Do, 2010), and assuming random mating. The method is robust to equilibrium migration rates up to 10% at lower population sizes (Waples & England, 2011; Gilbert & Whitlock, 2015). We performed analyses both for the whole dataset and separately for each of the four protected areas. The software estimates confidence intervals (CIs), both parametric and based on jackknifing over individuals (Jones et al., 2016), which accounts for the fact that overlapping pairs of loci are being compared and implements a method to correct for possible biases due to missing data (Peel et al., 2013). In any case, for each of the five datasets, we also performed analyses removing loci with more than 10% missing data. When analyzing each dataset, and depending on its sample size, we used a minimum allele frequency (MAF), the Pcrit value, following the recommendations in Waples & Do (2010).

## 3. Results

### 3.1 Effective population size estimation and demographic history

#### 3.1.1 The PSMC (pairwise sequentially Markovian coalescent) and the IICR

The PSMC curves exhibited a series of increases and decreases with the last decrease starting around 200 kya, and the previous one starting at around one million years ago (Figure 2a). These curves can be interpreted in terms of changes in *N_e_*, assuming panmixia and no population structure, in terms of changes in connectivity under population structure (Steux et al., 2024) and constant size or as a combination of both types of changes. Altogether they suggest either very large populations in the past that have been significantly reducing for the last 200 ky or are the result of a metapopulation characterized by changes in connectivity with no obvious population decrease during that period. The results for the 10 bootstrap replicates for each individual were highly consistent (Supplementary material, Figure S5). One individual (sample T3_chimp) exhibited a PSMC curve that followed the shape of the other curves but was flatter and shifted downward and towards more recent times, a pattern that has been described for individuals having a lower coverage (Nadachowska et al., 2016, see Discussion).

#### 3.1.2 MSVAR analysis of microsatellite data

The analyses we conducted with the control datasets (unrelated and “random datasets”) did not provide results significantly different from the demographic scenarios obtained for the complete datasets of GB and the parks, hence suggesting that relatedness and sample size did not significantly affect the results.

From the whole dataset (N = 143 genotypes), MSVAR estimated a contemporary *N_e_* (*N_o_*) between 4,500 and 6,500 individuals, that resulted from a mild bottleneck starting about 44,500-70,000 years ago from a more ancient *N_e_* around 10,000-12,000 (*N_1_*) (Fig. 2b). To account for the possible effects of genetic structure, we also carried out analyses separately for each park (Fig. 2e to 2h). For CNP, the southernmost chimpanzee population of GB, MSVAR identified a stronger signal with estimates of *N_0_* between approximately 500 and 1,125 individuals, whereas *N_1_* estimates were between 10,000 to 12,000 breeding individuals, with limited overlap between the *N_0_* and *N_1_* median posterior distributions and a posterior distribution of N_0_/N_1_ that was consistently below zero (Fig. 2e, Supplementary material, Table S5). MSVAR estimated that this demographic decrease of the CNP chimpanzee population occurred around 5,000 and 12,500 years ago under the assumption of panmixia (Table 2).

**Table 2.**
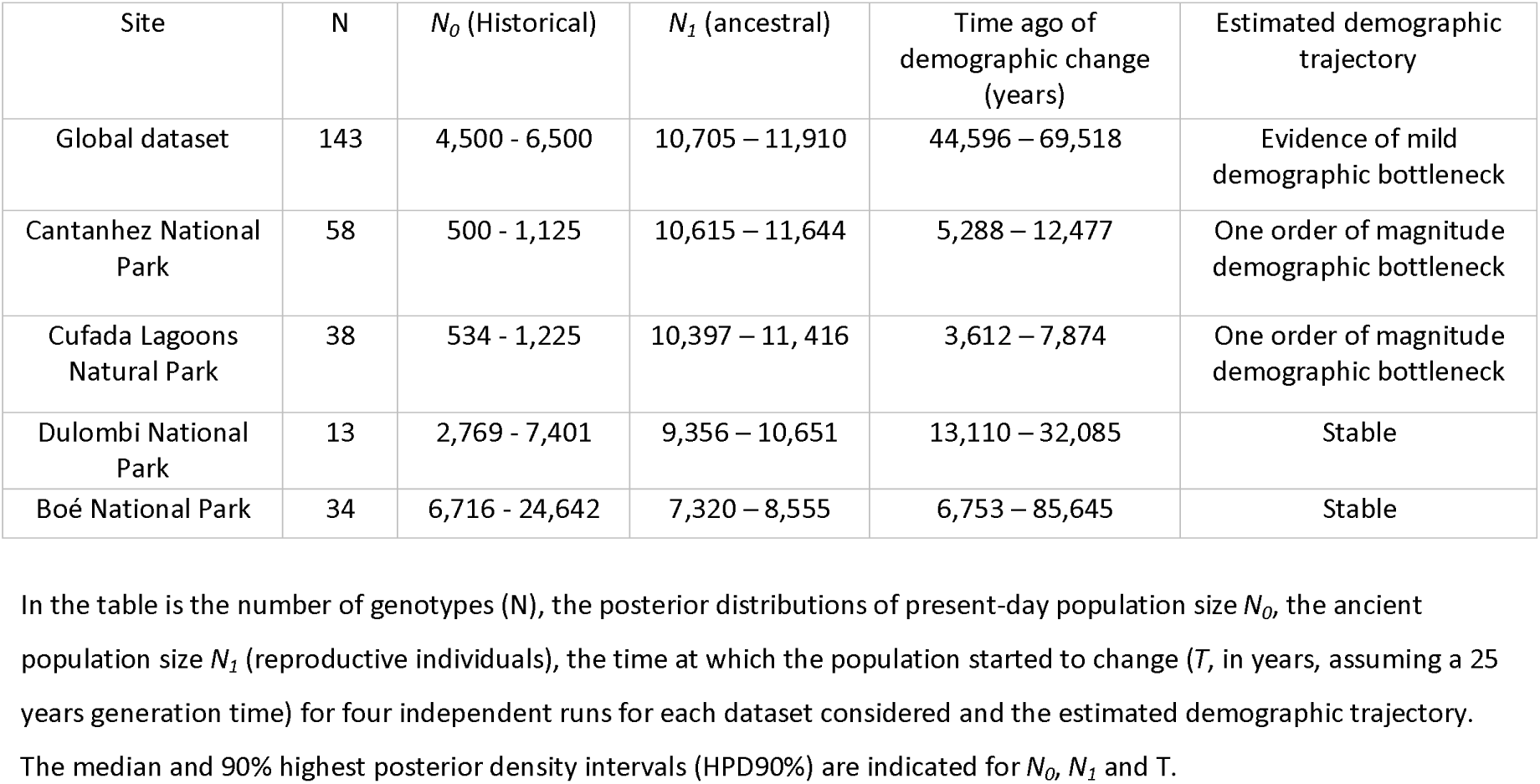
Estimation of long-term *N_e_* for the whole dataset and per park by employing the Bayesian likelihood-based approach implemented in MSVAR 1.3 (Storz & Beaumont 2002) and using microsatellite loci data.

| Site | N | $N_0$ (Historical) | $N_1$ (ancestral) | Time ago of demographic change (years) | Estimated demographic trajectory |
| --- | --- | --- | --- | --- | --- |
| Global dataset | 143 | 4,500 - 6,500 | 10,705 – 11,910 | 44,596 – 69,518 | Evidence of mild demographic bottleneck |
| Cantanhez National Park | 58 | 500 - 1,125 | 10,615 – 11,644 | 5,288 – 12,477 | One order of magnitude demographic bottleneck |
| Cufada Lagoons Natural Park | 38 | 534 - 1,225 | 10,397 – 11,416 | 3,612 – 7,874 | One order of magnitude demographic bottleneck |
| Dulombi National Park | 13 | 2,769 - 7,401 | 9,356 – 10,651 | 13,110 – 32,085 | Stable |
| Boé National Park | 34 | 6,716 - 24,642 | 7,320 – 8,555 | 6,753 – 85,645 | Stable |

In the table is the number of genotypes (N), the posterior distributions of present-day population size $N_0$ , the ancient population size $N_1$ (reproductive individuals), the time at which the population started to change ( $T$ , in years, assuming a 25 years generation time) for four independent runs for each dataset considered and the estimated demographic trajectory. The median and 90% highest posterior density intervals (HPD90%) are indicated for $N_0$ , $N_1$ and $T$ .

We found a similar demographic scenario for the CLNP chimpanzee population, which, like the CNP population, is also located in the coastal area and is geographically the closest to CNP (Fig. 2f). The CLNP chimpanzee population was also found to have undergone a one-order-of-magnitude bottleneck, with very similar values of *N_0_* and *N_1_* to the inferred scenario at CNP (Table 2). The putative difference between the demographic histories of the two coastal populations is the fact that the demographic decrease may have occurred later in CLNP than in CNP (CLNP: 3,612 – 7,874 years ago), but the inferred posterior distributions of T for both parks still overlap considerably (Table 2, Fig. 2e and Fig. 2f). The bottleneck signal of both populations is also confirmed by the *N_0_/N_1_*, which in both cases is below 0, across the four simulated scenarios (Fig. 2e and Fig. 2f).

As for the chimpanzee populations from the eastern region, at DNP and BNP, we found different demographic histories (Fig. 2g and Fig. 2h). Not only are the current *N_e_* much larger but there is also no clear signal of demographic change for any of these regions. The population from DNP is estimated to have a *N_0_* of between 2,769 and 7,401 individuals, with a slightly larger estimated *N_1_* (Table 2). However, the variation in *N_0_* estimates across the four different simulated scenarios does not allow for a clear signal of population size change. This absence of a clear bottleneck signal is also supported by the variation of the *N_0_*/*N_1_* estimates and its overlap with zero (meaning no population size change), which is also translated into weak convergence of the *T* posterior distributions (Fig. 2g). For the easternmost chimpanzee population sampled in BNP, the inferred posterior distributions of *N_0_* and *N_1_* are very consistent, which indicates a large and historically stable population in this park (Fig. 2h). The estimated values for the *N_0_* indicate a population with a current effective population size above 6,500 individuals (Table 2). The estimated posterior distributions of *N_1_* extensively overlap with the *N_0_* estimates, as it is also clear from Fig. 2h suggesting no population size changes. The scenario of a large and historically stable population is further supported by the *N_0_*/*N_1_* estimates consistently overlapping with zero (Fig 2h).

#### 3.1.3 Linkage disequilibrium-based estimation of current Ne and recent demographic history from genomic data

GONE estimated a contemporary *N_e_* around 300-350 (point estimate of 344 for the previous generation, 95% CI = [116, 580]). Regarding demographic history over the last 100 generations, analyses in GONE suggest that after an increase in *N_e_* over 50 generations, the effective population size gradually declined over the last 50 generations up to present times, following an almost symmetrical pattern (Fig. 2c). The *N_e_* decrease was of almost one order of magnitude.

#### 3.1.4 Linkage disequilibrium estimation of contemporary Ne from microsatellite data

Estimates of contemporary *N_e_* and respective 95% confidence intervals, obtained with NeEstimator from the microsatellite data, both for the global dataset and separately for each of the four parks, are presented in Table 3. The *N_e_* point estimates for the global dataset were around 150-190 individuals, with the 95% CIs spreading around 80-550 individuals.

Point estimates for CLNP and BNP were of similar magnitude, respectively at about 160-230 and 130-150 individuals, but the 95% CIs had infinite upper bounds. This can happen if *N_e_* is large and/or if the data has limited information (*e.g.,* insufficient sample size; reduced number or not very polymorphic loci) (Marandel et al., 2019). The infinite upper bound implies that it is not possible to reject the null hypothesis that LD can be explained entirely by sampling error (Waples & Do, 2010). Still, the finite lower bound provides useful information about the minimum limit of *N_e_* (Waples & Do, 2010). The *N_e_* point estimates for the CNP (54-79 individuals) were substantially lower, but with 95% CIs overlapping with the previous ones. On the other hand, the point estimates for DNP, at 8-12 individuals, were approximately an order of magnitude smaller than for the other datasets and with parametric 95% CIs not overlapping with the other parks; yet, the jackknife CIs had infinite upper bounds. The small sample size from the DNP may contribute to an underestimation of *N_e_*. This underestimation will tend to be smaller if the true *N_e_* is not very large (e.g., ≤ 100) and will be greater the larger the true *N_e_* (Waples & Do, 2010).

## 4. Discussion

In this work, we used several methods that aim to estimate either a single effective population size or possible changes in *N_e_* over different temporal scales, using samples obtained over different spatial scales. The genetic diversity of the critically endangered western chimpanzee in GB was estimated using different types of data (microsatellites and WGS SNPs) and the methods applied were thus different. We estimated the demographic trajectories of western chimpanzees’ representative of the whole country and, separately, for four geographic populations inhabiting protected areas in the south of the country, which are considered relevant areas given the global conservation of the subspecies.

### 4.1. Estimation of *N_e_* over different temporal scales

As noted in Luikart et al. (2010) review and in Box 1 in Ryman et al. (2019) one can consider, as a simple approximation, the idea that different *N_e_* estimators should be interpreted by using different time frames (Luikart et al., 2010, Wang, 2005). Some estimates would correspond to the “ancient time” (from hundreds to tens of thousands of generations) whereas others would correspond to the recent or contemporary *N_e_* (from a few to tens or hundreds of generations). Typically, MSVAR (microsatellite data) and PSMC (single genome data) provide information about the former time period whereas NeEstimator (microsatellite data) and GONE (WGS data) provide estimates that are mainly about the recent past. The latter type of methods assume that the long-term *N_e_* can be to some extent neglected regarding some properties of the genetic data such as the LD pattern (at least among some markers) or the variation of allele frequencies in the last few generations. We note that MSVAR also provides an estimate of contemporary *N_e_* but as part of a demographic model of size change, and that GONE also integrates the contemporary *N_e_* in a trajectory of *N_e_* change.

The contemporary *N_e_* estimates are considered the most relevant to assess extinction risk because they reflect ongoing or recent demographic or reproductive processes whereas the historical *N_e_* refers to the genetic or demographic processes over much longer periods (Luikart et al., 2010; Santos-del-Blanco et al., 2022). We argue that it is the combination of these estimates that should inform on the best conservation decisions and measures (Fig. 2).

Furthermore, although the concept of *N_e_* has been presented as analogous to the census size (*N_c_*), decades of research have shown repeatedly that *N_e_* and *N_c_* are not only distinct but may have nearly opposite trends under some models. We want to stress here that *N_e_* is only informative about some property of the genetic data, and different properties may have different temporal dynamics which are themselves possibly different from the *N_c_* dynamics (see Chikhi et al., 2018, Vishwakarma et al., 2024, Wakeley, 1999). *N_c_* may also be very disconnected from any *N_e_* estimate because *N_c_* informs us about the current individuals living in the environment of interest and are thus directly affecting and affected by major ecological processes, such as predation, competition or density dependence (Waples, 2022). Different *N_e_* values will depend on the probability that individuals have to contribute genes to the next generation but also on how populations may be connected to each other, as this will influence coalescence times (Beaumont, 2004, Chikhi et al., 2010, Wakeley, 1999). Contemporary *N_e_* will be influenced by recent fluctuations in population size, variance in reproductive success among individuals, unequal sex ratio and overlapping generations (Hoban et al., 2020, Waples, 2022). Over longer time periods (“historical *N_e_*”), population structure and changes in connectivity may generate very contradictory results. For instance, Wakeley (1999) showed that a structured population where all the demes increase in size and between which, gene flow increases at the same time, may exhibit a signal of decrease in *N_e_* in stark contrast with the increase of the actual population size. A similar phenomenon was described by Mazet et al. (2016) on the PSMC method (see also Parreira et al., this Special issue on the effect of population structure on different *N_e_* estimation methods).

The PSMC plots that we obtained with the five individuals from GB exhibited a similar trajectory to those obtained by Prado-Martinez et al. (2013) for individuals from the same subspecies (*P. t. verus*) but from other locations. The authors estimated the peak of effective population size at ∼150 kyr, which is similar to the time of the highest *N_e_* estimated here (around 200 kyr). The difference in these values could be due to the fact that we used the chimpanzee reference genome, whereas Prado-Martinez et al. (2013) used the human genome as a reference. However, these differences are minimal, probably because the divergence between *Homo sapiens* and *P. troglodytes* is on the order of 1% (see Henrique et al., 2024 and references therein for the effect of divergence of the reference genome). Beyond these important technical issues, the PSMC curves must be interpreted with care as they could indicate changes in *N_e_* through time, changes in connectivity without change in *N_e_* (Steux et al., 2024) or, more likely, a combination of both changes in *N_e_* and connectivity. One particularly striking result was that individual T_3 exhibited a PSMC lower than the other four individuals, and shifted towards the left (more recent times). Several studies have found that this kind of shift can be observed when the coverage is reduced (Nadachowska et al., 2016 on *Ficedula* sp. flycatchers and Henrique et al., 2024 on *Microcebus* sp. mouse lemurs and several other primate species). However, individual T_3 has a higher rather than smaller coverage than the other individuals (T_3 32 X vs. 17 X for Bella, Bo, Emília and Simão, on average). This result is thus puzzling, and it cannot be explained without further investigation. One possible explanation is that the difference in coverage between samples may have resulted in variable power to call heterozygous sites, which could have resulted in the slightly distinct demographic histories estimated for the five individuals (Frantz et al., 2013).

The PSMC method has been used on many endangered species since it only requires one genome sequence and is thus adapted for endangered species for which large genomic data sets are difficult to obtain. This is the case of many endangered primates, including chimpanzees (Prado-Martinez et al., 2013), gibbons (Hylobatidae *sp*., Carbone et al., 2014), mouse lemurs (Teixeira et al., 2021), and sifakas (Propithecus *sp*., Guevara et al., 2022). Implicitly, what the PSMC recovers is the distribution of coalescence times along the genome. When two non-recombining fragments are identified by the PSMC and differ by many heterozygous sites, this suggests that their MRCA (most recent common ancestor) is likely very old. The opposite is more likely if there are few or no heterozygous sites. The distribution of these implicit coalescence times can be interpreted as the result of population size changes under the assumption of panmixia and in the absence of significant population structure in the species of interest. The PSMC can thus be interpreted as a series of changes in *N_e_*, and this is the most common interpretation.

However, in the last 25 years there has been an increasing recognition that population structure can generate spurious signatures of population size change (Beaumont, 2004, Chikhi et al., 2010, Wakeley, 1999). In the specific case of the PSMC method, Mazet et al. (2016) showed that it is in fact impossible to determine whether a particular PSMC plot is the result of real change in *N_e_* or of a more complex model of population structure with changes in connectivity, without any change in population size. In the latter case, it thus becomes impossible to actually make statements regarding changes in *N_e_* from changes observed in the PSMC curve alone. Mazet et al. (2016) introduced the concept of IICR (inverse instantaneous coalescence rate) and noted that the PSMC method in fact infers the IICR, not *N_e_*. The IICR will be identical to *N_e_* under panmictic models without population structure but very different from any *N_e_* changes as soon as there is population structure. Altogether, this suggests that for species like chimpanzees that are known to be structured (Funfstuck et al., 2015, Lester et al., 2021), signals of population size changes inferred from methods assuming panmixia (PSMC, MSVAR, Bottleneck, StairwayPlot, GONE, etc.) must be interpreted with caution (Steux et al., 2024). However, whether one considers population structure or panmixia, our results suggest that the populations sampled were part of a metapopulation that may have been very large and included all the regions sampled in this paper. We will come back to this later.

The analyses using the MSVAR method and microsatellite loci data suggested that chimpanzees from GB have undergone a mild demographic decrease starting around 40,000 years ago (when we used the global dataset). However, for the analyses of the different parks, we inferred contrasting histories for inland (eastern, DNP and BNP) and coastal (western, CLNP and CNP) populations with either no changes or recent and minor *N_e_* changes. Also, the estimates for *N_1_* were very similar for the different parks with ∼10,000 individuals. This may indicate that the populations at the four parks were connected in the past and part of one metapopulation (itself probably connected to other regions outside GB), in agreement with our interpretation of the PSMC curves.

By contrast, the estimates of *N_0_* were much smaller, indicating either that population connectivity changed towards more recent times, or that these values correspond to the local deme size, as expected under the coalescent theory of structured models. Whichever interpretation one may favor, our results appear to suggest that, whereas the inland eastern populations remained large/connected with other regions (*e.g.*, transnational populations, given the geographic proximity with the Republic of Guinea), the western coastal populations appear more isolated, a process that may have started thousands of years ago at the scale of the country. We have to be cautious with these figures as the timing of size change inferred by MSVAR may not correspond to any particular timing of change in gene flow, since models without changes in connectivity would also generate signals of bottleneck. Despite these cautionary remarks, we believe that the CNP and CLNP may have become separated from the metapopulation, but could have remained connected to each other, forming a smaller sub-population not exceeding 1,200 breeding individuals. This result also suggests that chimpanzees may have still been able to disperse between these two parks.

In a recent study, Fontsere et al. (2022) analyzed chromosome 21 genome-wide data across the species and subspecies distribution and inferred a large exchange of migrants during the last ∼800 years (range 117 to 2,200 years) for the populations located in the northern range of the distribution, which includes Senegal, Mali, northern Guinea and GB (using samples from BNP only). Thus, no signs of long-term isolation were detected for GB. In another study, Heinicke et al. (2019a) investigated the existence of subpopulations across the *P. t. verus* distribution based on field survey data and spatial modeling tools. According to these authors, one large subpopulation (> 33,000 individuals, approximately 50% of the total population size) was predicted at the northern range of the subspecies, in areas characterized by savanna-mosaic habitats and extending across the Fouta Djallon highland region and the neighboring areas of Senegal and GB (including the four parks of our study), up to Mali and Sierra Leone. Thus, these studies could explain why we inferred large *N_1_* estimates of ∼10,000 reproductive individuals (compared to *N_0_*). These estimates could correspond to the whole GB population, possibly reflecting an historical connection of the GB population to a large metapopulation centered in the Fouta Djallon highland region. In a more recent study, Steux et al. (2024) suggested that patterns of genomic variation as observed in the PSMC curves could be modeled as part of a metapopulation of small demes characterized by periods of changing connectivity. They estimated that, in comparison to other *P. troglodytes* subspecies, *P. t. verus* population was characterized by smaller demes, which could explain the lower nucleotide diversity observed in western chimpanzees. Their study however focused on the rather ancient past (older than ca. 50 kyr) due to the uncertainties on the PSMC curves they analyzed in the recent past. Other genetic studies based on microsatellite loci and a fragment of the mitochondrial DNA D-loop region, carried out in GB, do not indicate a strong population structure (Borges, 2017; Sá, 2013), which suggest that the chimpanzees are able to disperse between parks. The change in connectivity between inland areas (DNP-BNP) and coastal (CNP-CLNP) within GB cannot be directly inferred from our analyses. We must thus be careful in result interpretations. However, our analyses do identify a large ancestral population that might have been fragmented as a consequence of environmental changes that occurred around 10,000 years ago, including climate instability in West Africa and the more recent increased human impact resulting from the development of agriculture. The Younger-Dryas Holocene transition marks a very abrupt transition between the African Humid Period (14,500 - 6,000 years ago, characterized by the expansion of forests and lakes across the Sahara region), and a sequence of time periods characterized by arid conditions towards the late Holocene (Gasse & Van Campo, 1994). This climate instability, with a succession of warm and cooling events in West Africa, impacted the extension of forest cover (deMenocal et al., 2000) and, most likely, the size and connectivity of the populations of forest-dwelling fauna, as is the case of the western chimpanzee.

The GONE analyses suggested a surprising growth of *N_e_* between 2,500 and 1,250 years ago (*i.e.*, in generations 100-50) followed by a short stationary period and a more recent symmetrical decrease. This pattern may be an artifact of the method, due to the small number of individuals analyzed (five chimpanzees). For example, Beichman et al. (2018) illustrated the general difficulties of demographic history methods in inferring demographic events over the past hundred generations using whole-genome data for fewer than 10 individuals. Reid & Pinsky (2022) also emphasized the importance of large sample sizes for different demographic history methods regarding power and precision to detect and quantify population declines in the last 100 generations, with GONE being no exception to a sharp deterioration in performance when sample sizes are small. Similar and even larger humps have also been described by Santiago et al. (2020) when they simulated data from a simple structured model (panel F of their Figure 2), but at this stage one should be cautious to interpret these results until larger sample sizes are analyzed.

The result that however appears to be consistent with the other methods used in this study, is that a gradual decline of *N_e_* took place during the last 1,250 years (*i.e.*, the last 50 chimpanzee generations) towards the present, with the whole population reaching a contemporary *N_e_* around 300-350 breeding individuals, estimates that are qualitatively in agreement with those of NeEstimator and microsatellite data (*N_e_* of 107-549 or 138-294, respectively).

### 4.2 Implications for conservation management of the western chimpanzee in Guinea-Bissau

*N_e_* should be seen as a key parameter in conservation management and genetic biodiversity monitoring because it is supposed to provide information on population viability, on inbreeding depression risk, on population isolation, and on effectiveness of selective processes and adaptation in relation to drift (Charlesworth, 2009, Hoban et al., 2022). In populations with small *N_e_*, genetic diversity is lost at a faster rate over time, and the random fluctuations in allele frequency by drift can neutralize the effect of natural selection and increase the probability of fixation of deleterious mutations, potentially leading the population to inbreeding depression (Charlesworth 2009, Hoban et al., 2022, Newman & Pilson, 1997).

Franklin (1980) proposed the thresholds of 50/500 for a minimum effective size required for a viable population in the short and long term, respectively. This recommendation became an established rule of thumb in conservation biology and has been proposed as a genetic indicator to assess progress towards global conservation targets (Frankham et al., 2014, Hoban et al., 2020). The 50 short-term rule refers to an effective size quantifying the rate of inbreeding (*N_eI_*). The minimum *N_eI_* of 50 individuals is thought to be enough to prevent the rapid inbreeding of the population (*i.e.,* 1% per generation), which could lead to excessive homozygosity for deleterious recessive alleles and reduced fitness by inbreeding depression. The 500 long-term rule refers to the effective size related to the loss of additive genetic variation (*N_eAV_*). This threshold defines the *N_e_* above which a population should retain enough evolutionary potential to adapt to new selective forces (*i.e.,* future environmental conditions, Jamieson & Allendorf, 2012). More recently, these numbers have been doubled, with 100 individuals being presented as more adequate to prevent inbreeding depression over five generations for wild populations (*i.e.*, limiting to 10% the loss in total fitness) and 1,000 individuals as necessary to protect evolutionary potential in the long-term (Frankham et al., 2014), particularly when the specieś reproductive rates are low (Perez-Pereira et al., 2022). When a population is detected to have a small or declining *N_e_*, managers and conservationists should be called to investigate the most likely causes and to reverse the demographic trajectory (Wang et al., 2016). This is why estimating *N_e_* is increasingly recognized as central to conservation programs.

In this work, we used different approaches that estimated historical and contemporary *N_e_*. While the estimates by the different methods could differ, results were consistent in suggesting that current *N_e_* is below 500 breeding individuals for the GB chimpanzee’s population. For instance, GONE and NeEstimator suggested values that were typically between 100 and 500, even if some estimates could have larger upper bounds, most likely due to small sample sizes (see Results section). These results were qualitatively in agreement with the *N_o_* estimates obtained with MSVAR even if the latter were higher (between 500 and 1,000). While no *N_e_* estimate should be taken at face value, the fact that these estimates were qualitatively similar when using different methods making different assumptions and using different types of genetic markers, suggests that the contemporary demographic dynamics of chimpanzees in GB is driven by small and isolated populations that derive from, what used to be, a very large metapopulation. Furthermore, these results confirm the selection of the coastal areas of GB and Republic of Guinea by Kormos & Boesch (2003) as priority regions for conservation of the subspecies and are aligned with estimates of density that points to a small population in CLNP (Carvalho et al., 2013), highlighting the critical conservation situation of these populations.

Small populations (*i.e*., < 500 breeding individuals) may go extinct through a phenomenon referred as extinction vortex (Gilpin & Soule, 1986), in which genetic and demographic issues interact synergistically to decrease genetic diversity and cause population growth rate to drop due to the reduction of the mean fitness. This results in further decreases in genetic diversity and promotes the subsequent processes to happen in a cascade, until the extinction of the population. In the case of the western chimpanzees in GB, the main conservation threats have been identified and characterized to some extent. Natural habitats have generally been converted into subsistence crop plantations, such as rice (*Oryza* spp.) or cassava (*Manihot esculenta*), and cashew (*Anacardium occidentale*) monoculture agroforests at least since the last two decades (Hockings & Sousa, 2013, Temudo & Abrantes, 2014). Additionally, the construction of roads and infrastructures increased the accessibility to remote areas and promoted more encounters with humans, which may have increased chimpanzees’ mortality. Although it was found that chimpanzees can cross and use human-altered habitats to some degree, namely sharing the use of forested and village areas with local communities (*e.g*., in CNP, Bersacola et al., 2021), extensive habitat loss and conversion into crop fields and villages are expected to reduce connectivity between populations and diminish population size rapidly (Torres et al., 2010). In CNP for instance, it was estimated a loss of 11% of suitable habitat and the death of between 157 and 1,103 individuals in the population for the period between 1986 and 2003 (Torres et al., 2010), which corresponds to less than one chimpanzee generation. Moreover, as reported by Stiles (2023) and as illustrated here (with the four blood samples of confiscated individuals), hunting for live individuals to supply the national and international illegal pet trade occurs in the country (Ferreira da Silva & Regalla, in review, Stiles 2023). Quantitative data on the number of traded chimpanzees originating from the GB in international trade routes is missing (Clough & Channing, 2018; Stiles et al., 2013; Stiles, 2023). However, given the ease of detecting chimpanzees in private houses and hotels (*e.g.*, 18 individuals between 2006 and 2022 found by chance, in Ferreira da Silva & Regalla in review) and considering that five to 10 adults can be killed to harvest only one infant chimpanzee (Teleki, 1980), it can be suggested that hunting to supply the trade of live individuals may have contributed to a reduction of the population size and consequently of the *N_e_*. Furthermore, as conservation threats tend to act synergistically at the local and regional scale - habitat fragmentation increases accessibility to natural habitats by local communities, which in turn, increase poaching, negative interactions with farmers and diseases transmission (Humle et al., 2016), the negative impacts of human-derived activities on the population effective size may be larger than each threat is individually.

Specifically, chimpanzees inhabiting CNP and CLNP - the two populations identified by this study at a high risk of extinction - are currently negatively impacted by habitat loss and fragmentation, and by retaliatory killing by farmers during crop-raiding (Hockings & Sousa, 2013). Chimpanzees in CNP may also be subjected to higher reproductive isolation since the Park is in a peninsula surrounded by two permanent water bodies which are insurmountable by chimpanzees, and suitable habitat for the subspecies was considerably lost in northwestern areas, where the isthmus connects the peninsula to the mainland (*i.e*., Bantael Sila, Cumbijã and Guiledge villages, Torres et al., 2010). These coastal areas have been considered important to maintain gene flow with GB mainland for another primate species (*e.g.,* Guinea baboons, *Papio papio*, Ferreira da Silva et al., 2014, Ferreira da Silva et al., in press). Other forest-dwelling and hunted primate species at CNP may have gone through a decline in *N_e_* of a similar magnitude. Minhós et al. (2016) found a pattern of *N_e_* decrease for co-distributed populations of colobus monkeys inhabiting CNP (*Piliocolobis badius temminckii* and *Colobus polykomos*) at a similar time period as estimated here for CNP chimpanzees (*i.e.*, ca. 10,000 to 3,000 years). The *N_e_* for colobus monkeys was estimated using alike models and genetic markers (*i.e*., MSVAR runs and microsatellite loci). The fact that a similar demographic event was observed for other co-distributed species in CNP despite different socio-ecology features, strengthens our interpretation that CNP chimpanzees experienced low *N_e_* in recent times. Moreover, several chimpanzee communities across CNP show symptoms of leprosy (caused by *Mycobacterium leprae*, Hockings et al., 2021), which is likely to have negative consequences for the longevity and reproductive success of individuals, but so far unknown. On the other hand, In the southern region of CLNP, the construction of a large road and a thermoelectric plant with respective electricity transmission lines, led to the loss of one of the best-preserved forest patches of the park (Catarino, 2019) and increased the accessibility to areas used by chimpanzees for nesting (Carvalho et al., 2013). Cases of live chimpanzee captures have been recorded in both parks, which most likely implies adult mortality (Ferreira da Silva & Regalla in review). Furthermore, CLNP is bordered in the east by the main road connecting the south of the country to the capital city - Bissau and, as demonstrated in this study, wildlife-vehicle collisions do happen. Our results suggest that CLNP and CNP are at high risk of extinction and the impact of human-derived activities potentially threatening the two chimpanzee populations should be investigated.

Our study provides the first estimates of *N_e_* for DNP. We estimated an historical *N_e_* of 2,769 - 7,401 breeding individuals using MSVAR but could not detect a strong departure from mutation-drift equilibrium. This result was at odds with the very low contemporary *N_e_* of 6-30 obtained using NeEstimator. DNP is located at the northern margin of the Corubal River and is currently at one edge of the subspecies distribution (see Fig. 1b). Our estimates of a stable demographic trajectory and a large historical *N_e_* suggest that DNP was at some point in time connected to other large chimpanzee populations. Connectivity to populations located in the south of the Corubal River (*e.g.*, BNP) should be more reduced at present times because the current width of the water body, which can surpass one kilometer in some sites, may be a significant barrier to primate’s gene-flow (*e.g*., the green monkey, *Chlorocebus sabaeus*, Colmonero-Costeira et al., in review, the Guinea baboon, *Papio papio*, Ferreira da Silva et al., in press). Nevertheless, the configuration and discharge of the Corubal River may have been different in the past (*e.g*., the mouth may have been in the southwest part of the country, located by the Rio Grande de Buba channel area, Alves 2007), which could have allowed chimpanzees to cross to what is the south margin. The small contemporary *N_e_* may be explained by the fact that present environmental conditions do not support a large population of chimpanzees. DNP is located at the edge of the distribution of the subspecies and has low density of villages and other human infrastructures. This area is mostly dominated by woodlands and savannah woodland formations (Catarino et al., 2008), and found to be of low habitat suitability for chimpanzees by modeling exercises (<100, range 0-1,000, in Figure S2.2. Carvalho et al., 2021), which could be either related to environmental conditions or a small sample size (J. Carvalho, personal communication). During field work, chimpanzees were mostly detected (and fecal samples collected) in greater proximity to gallery forests along smaller streams or next to the Corubal River (Fig. 1b), and we observed that the subspecies was not widely distributed in the park area, as in CNP, for instance (Sá, 2013). Chimpanzees at Fongoli in Senegal, inhabiting a similar open, savanna-woodland environment, do not suffer from nutritional stress but display physiological stress from dehydration and heat, which does not seem to be behaviorally compensated (from sitting for longer periods in the shade or using pools or caves for instance, Wessling et al., 2018). Such adverse environmental conditions may be determinant for constraining the distribution at the biogeographical range limits of the subspecies (Wessling et al., 2018) and similarly, limiting the size of the population at DNP.

By contrast, our estimates of historical *N_e_* of the population of chimpanzees inhabiting the BNP (MSVAR *N_0_*: 6,716 - 24,642 breeding individuals) are large and confirm the classification of the area as stable or of high-density (Heinicke et al., 2019b). Boé population has been included in a previous population genomic study, using samples collected across the subspecies range (Fontesere et al., 2022), which estimated high and recent connectivity (for the last ∼780 years, range 117-2,200 years) between communities at the northern range (localities in the Republic of Guinea and south of Senegal, Fig. 3 in Fontesere et al., 2022). Boé was found to be genetically closer to samples collected in southern Senegal (Fontesere et al., 2022), possibly due to long-term connectivity between the two neighboring populations. Present-day high density of chimpanzees in the Boé region has been justified by i) remoteness of the area and difficult access, ii) rare hunting of chimpanzee to comply with religious taboos, iii) high habitat suitability for chimpanzees, and iv) slow habitat loss and conversion, and in a large area, habitats are undisturbed (Binczik et al., 2019; Carvalho et al., 2021, van Laar 2010). Although DNP and BNP are closely located and share similar environmental conditions, within the Boé region there is a wide network of rivers and waterbodies surrounded by relatively well-preserved gallery forests, which are used by chimpanzees to nest and feed (Binczik et al., 2019). Our results suggest that Boé is a stronghold for the chimpanzee population in GB. The effective protection and restoration of the natural habitats in ecological corridors connecting BNP and the remaining parks located south of the Corubal River (Fig. 1b) could be beneficial to promote dispersal, potentially increasing gene flow and improving the probability for long-term persistence of chimpanzees in coastal areas of GB.

### 4.3 Implications for the estimation of *N_e_* of wild populations of primates

The most common approaches to estimate the effective size of real populations are based on its genetic properties (Luikart et al., 2010). However, obtaining genetic information of threatened species can be challenging. The main issue is related to attaining high quality DNA, which is usually extracted from fresh blood and tissue samples. Species of conservation concern are frequently found in low densities in the wild and commonly live in inaccessible areas and in habitats of low visibility. Hence, it is difficult to trap or handle individuals (Beja-Pereira et al., 2009). Moreover, invasive methods are considered unethical as the contact with humans to retrieve blood or tissue samples increases the risk for disease transmission (Beja-Pereira et al., 2009). Thus, invasive sampling of wild-born individuals for threatened species is typically opportunistic and carried out for a few individuals, for instance during veterinarian interventions or post-mortem (*e.g*., Prado-Martinez et al., 2013). Such procedures can take several years to complete (Xue et al., 2015), and sampling is usually geographically restricted, and not representative, of the species distribution and variability. In primates, genomic data used to infer parameters of interest for conservation, has been obtained from high quality DNA collected from individuals in zoos or rescuing centers (*e.g.*, Rogers et al., 2019 but see Fontsere et al., 2022 who obtained genomic diversity estimates of wild chimpanzee populations from 828 non-invasively fecal samples). Yet, specific environmental conditions or breeding practices that inadvertently reduce natural selection pressures or increase inbreeding (Christie et al., 2012) can lead to wrong or limited inferences of demographic parameters. Also, small and spatially restricted sampling can introduce bias in contemporary *N*_e_ estimates (*e.g*., Santos-del-Blanco et al., 2021) and geographic-broad genetic data, such as the ones obtained using non-invasive fecal samples, are preferable.

Here, we show that the estimated values of *N_e_* using genomic data and more classic genetic markers, like microsatellite loci obtained using non-invasive fecal samples, are largely concordant, although we found that median *N_e_* estimates produced by SNP data were higher than estimates generated using microsatellite data. This pattern was also reported by Clarke et al. (2024) meta-analysis. Our study reinforces that datasets generated with traditional genetic markers, such as legacy or baseline microsatellite loci datasets for local populations, are of great value; these can be used to estimate parameters relevant to inform conservation management in species for which obtaining genomic data is not straightforward, or in studies carried out in countries with limited access to sequencing units, funding and trained researchers in genomic data (Bertola et al., 2024). Moreover, our study shows that the combination of different molecular markers and analytical methods can be a useful strategy to overcome the limitations of obtaining high quality DNA from wild threatened populations, to investigate species evolutionary history in time and space, and to integrate genetic information in conservation management decisions at local and regional scales.

## Acknowledgments

We dedicate this chapter to the memory of Michael W Bruford - our mentor, colleague and friend. His enthusiasm, guidance, and support were key to the success of this work and to advance the knowledge on conservation genetics of Guinea-Bissau primates. We would like to acknowledge the Guinea-Bissau governmental agency Instituto de Biodiversidade e Áreas Protegidas (IBAP), namely to the former director Dr. Alfredo Silva and Dr. Justino Biai, and to the directors of protected areas and staff members - Dr. Abilio Said, Dr. Augusto Cá, Dr. Joãozinho Mané and Dr. Sadjo Danfa, for fieldwork and sampling permits and to Abel Vieira, Iaia Cassama, Benjamin Indeque, Braima Bemba Canté for the support in fieldwork logistics. We acknowledge the Direcção Geral de Florestas e Fauna (DGFF) and CITES focal person in GB for samples exportation permits; to the research assistants and guides Sadjo Camará, Mamadu Soares, Mamadu Turé, Idrissa Camará; to the NGO CHIMBO for logistical support for carrying out fieldwork in the Boé region; to I. Espinosa and H. Foito for logistical support in Bissau. We thank Dr. Pedro Melo (vetnatura) and the embassy of the European Union in Bissau for the help in collecting blood samples from chimpanzees. We are grateful to Lara Almeida for facilitating the blood samples of the chimpanzee Simão. This research was funded by Fundação para a Ciência e Tecnologia through the project PRIMATOMICS (PTDC/IVC-ANT/3058/2014), and by funders of the PRIMACTION project (the Born Free Foundation, Chester Zoo Conservation Fund, Primate Conservation Incorporated, Mohamed Bin Zayed (Project 232533027) and by sponsorship by the following Portuguese private companies - CAROSI, Cápsulas do Norte, Camarc, JA-Rolhas e Cápsulas). MJFS worked under an FCT contract (https://doi.org/10.54499/CEECIND/01937/2017/CP1423/CT0010). I.C.C., F.B. and I.P were supported by FCT-doctoral fellowships (ICC: https://doi.org/10.54499/SFRH/BD/146509/2019, F.B.:https://doi.org/10.54499/2020.05839. BD; I.P.: https://doi.org/10.54499/SFRH/BD/118444/2016 with https://doi.org/10.54499/COVID/BD/151758/2021. M.C. was supported by a postdoctoral research associate contract (BBSRC, BB/R015260/1). R.M.S. was funded under: https://doi.org/10.54499/DL57/2016/CP1456/CT0002 and UID/00713/2020. C.R.F. thanks the support of CE3C through an assistant researcher contract (FCiência.ID contract #366) and FCT for Portuguese National Funds attributed to CE3C within the projects UIDB/00329/2020, UIDP/00329/2020, and LA/P/0121/2020, and FPUL for a contract of invited assistant professor

## Ethics

The manuscript has not been submitted elsewhere. The research complied with ethical guidelines, rules and protocols approved by IBAP and CIBIO-InBIO and adhered to the legal requirements of Guinea-Bissau (GB) and Portugal. All except five samples were obtained non-invasively from unidentified individuals without manipulation or perturbation of their daily behavior. Invasive samples (tissue and blood) were collected opportunistically from animals already deceased (tissue) or collected during health check to individuals living in captivity, by a certified veterinarian (Dr. P. Melo). The blood collection was approved by IBAP. GB CITES focal point (Direção Geral de Florestas e Fauna) authorized collection and exportation of blood and tissue samples. IBAP authorized collection of fecal samples in protected areas and transportation to Portugal. ICNF Portugal (Instituto para a Conservação da Natureza e Florestas) and DGV (Direção Geral de Veterinária) authorized importation of blood and fecal samples (Import Permits Simão 19PTLX00367, Tissue sample run-over individual N.°/ No 18PTLX005871 and Bo, Bella and Emilia blood samples 17-PT-LX00392/l).

## Informed Consent

Non-Applicable

## Data availability statement

The data that supports the findings of this study are available in Dryad Digital Repository.

## Funding statement

This research was funded by Fundação para a Ciência e Tecnologia (FCT) through the project PRIMATOMICS (PTDC/IVC-ANT/3058/2014), and by funders of the PRIMACTION project (the Born Free Foundation, Chester Zoo Conservation Fund, Primate Conservation Incorporated, Mohamed Bin Zayed (Project 232533027), and by sponsorship by the following Portuguese private - CAROSI, Cápsulas do Norte, Camarc, JA-Rolhas e Cápsulas). MJFS worked under an FCT contract (https://doi.org/10.54499/CEECIND/01937/2017/CP1423/CT0010). (ICC: https://doi.org/10.54499/SFRH/BD/146509/2019, F.B.: https://doi.org/10.54499/2020.05839.BD, I.P. https://doi.org/10.54499/SFRH/BD/118444/2016 with https://doi.org/10.54499/COVID/BD/151758/2021. Rui M. Sá was funded under: https://doi.org/10.54499/DL57/2016/CP1456/CT0002 and UID/00713/2020. M.C. was supported by a postdoctoral research associate contract (CryoArks project, Biotechnology and Biological Sciences Research Council, BB/R015260/1). C.R.F. was supported by an assistant researcher contract (FCiência.ID contract #366), FCT projects UIDB/00329/2020, and LA/P/0121/2020, and FPUL for a contract of invited assistant professor.

## Conflict of Interest

The authors declare that they have no known competing financial interests or personal relationships that could have appeared to influence the work reported in this paper.

## Compliance with International Conventions and Regulations on Biological Diversity and Endangered Species

We declare that this work complies with the Convention on Biological Diversity and the Convention on International Trade in Endangered Species of Wild Fauna and Flora (CBD and CITES). Within the CBD, we followed the Access to Benefit Sharing (ABS) guidelines. We give credit and equal access to benefits to the countries involved in the study (Guinea-Bissau, Portugal and the UK) and the respective academic institutions and scientists involved in the collection and analysis of data, who are co-authors to this work. We declare that we complied with CITES regulations and obtained export and import permits to move samples from Guinea-Bissau to Portugal for analyses, following CITES guidelines.

## Data Archiving Statement

Data for this study are available at Dryad Digital Repository: *to be completed after manuscript is accepted for publication*.

## Footnotes

3 See documentary about rescuing chimpanzees to Ol Pejeta sanctuary, Kenya, https://www.youtube.com/watch?v=GxXMk2UPvUM.

## Notes

### Competing Interest Statement

The authors have declared no competing interest.

