## Supplementary material for "Estimating the effective population size across space and time in the Critically Endangered western chimpanzee in Guinea-Bissau: challenges and implications for conservation management"

Ferreira da Silva et al

S1. Summary of DNA extraction protocol used in blood and tissue samples (adapted from Vallet et al., 2008)

The first day of DNA extraction starts with adding warmed CTAB 2% solution to the samples after which a first lysis was performed at 60 ºC and 700 rpm for two hours. A second lysis was performed using CTAB 10% and incubating the samples at 60 ºC for two hours at 700 rpm. Proteinase K and chloroform-isoamyl were added to the samples and a precipitation step using cold isopropanol at -20 ºC was carried-out overnight. The second day of DNA extraction included two washes using cold ethanol 70%. DNA was eluted in TE. A cleanup was performed using pre-washed magnetic SPRI beads. To perform a preliminary quality control of the extractions, a 2% agarose gel was run and DNA concentration was measured using Nanodrop. Laboratory procedures took place at the IGC, and extractions were carried out at a laminar flow hood in a Biosafety Level 2 dedicated room.

Table S2. DNA concentration measured using Nanodrop

| Sample | Concentration |
| --- | --- |
| Simão | 27,9 ng/µL |
| Bella | 53,9 ng/µL |
| Bo | 28,7 ng/µL |
| Emilia | 48,0 ng/µL |
| T-3 Chimp | 13,2 ng/µL |

Table S3. WGS data summary statistics for each sample

| Sample | Description | Coverage | Observed number of homozygous genotypes | Expected number of homozygous genotypes | Number of non-missing genotypes | Inbreeding coefficient (F) |
| --- | --- | --- | --- | --- | --- | --- |
| Bella_PT_GB | obtained from blood that was conserved in EDTA and room temperature until DNA extraction | 16.373 | 3469972 | 3.723e+06 | 5557472 | -0.1381 |
| Bo_PT_GB | obtained from blood that was conserved in EDTA and room temperature until DNA extraction | 16.683 | 3488163 | 3.723e+06 | 5557411 | -0.1282 |
| Emi_PT_GB | obtained from blood that was conserved in EDTA and room temperature until DNA extraction | 18.769 | 3529060 | 3.723e+06 | 5557167 | -0.1058 |
| Simao_PT_GB | obtained from blood that was conserved in EDTA and room temperature until DNA extraction | 17.154 | 3457122 | 3.723e+06 | 5557101 | -0.145 |
| T-3_Chimp | obtained from tissue (muscular) from a road-kill, found a few minutes after the accident. Sample was kept in 90% ethanol and room temperature until DNA extraction | 31.739 | 3425898 | 3.723e+06 | 5556748 | -0.1619 |

Table S4. Initial values and varying sets of priors and hyperpriors, and respective variance of the mean (in brackets), tested using MSVAR 1.3 to reflect assumptions of stable populations (*N_0_* = *N_1_*), population declines (bottleneck, *N_0_* < *N_1_*), or moderate expansion (*N_0_* > *N_1_*). N_0_ – Current effective population size, N_1_ – Past effective population size, T – time of the inferred demographic change.

|  | | **Model 1**  **Stable population** | **Model 2**  **Bottleneck** | **Model 3**  **Severe expansion** | **Model 4**  **Moderate expansion** |
| --- | --- | --- | --- | --- | --- |
| **Priors** | **l**og10(N_0_) | 4 1 | 3 1 | 5 1 | 5 1 |
|  | log(N_1_) | 4 1 | 5 1 | 3 1 | 4 1 |
|  | log(Ɵ) | -3.5 1 | -3.5 1 | -3.5 1 | -3.5 1 |
|  | log(T) | 5 1 | 5 1 | 4 1 | 4 1 |
| **Hyperpriors** | **l**og(N_0_) | 6 2 0 0.5 | 4 1.4 0 0.5 | 4 2 0 0.5 | 3 2 0 0.5 |
|  | log(N_1_) | 5 2 0 0.5 | 4 1.4 0 0.5 | 5 2 0 0.5 | 5 2 0 0.5 |
|  | log(Ɵ) | -3.5 0.25 0 0.5 | -3.5 0.25 0 0.5 | -3.5 0.25 0 0.5 | -3.5 0.25 0 0.5 |
|  | log(T) | 5 2 0 0.5 | 5 2 0 0.5 | 5 2 0 0.5 | 5 2 0 0.5 |

Table S5. Posterior distributions of the Median and 95% Highest Posterior density intervals (HPD; presented within brackets), calculated using BOA R package. Demographic parameters are as follows: *N_0_* - current effective population size; *N_1_* - past effective population size and T - time since the demographic change occurred in years.

|  |  | ***N_0_*** | ***N_1_*** | ***T*** |
| --- | --- | --- | --- | --- |
| ***GB*** | *Run 1* | 6 510 | 11 663 | 69 518 |
|  |  | (651 - 6,243,096) | (1060 - 302,134) | (31 - 140,637,132) |
|  | *Run 2* | 4 633 | 10 705 | 50 385 |
|  |  | (511 - 27,284) | (1435 - 95,675) | (106 - 56,728,330) |
|  | *Run 3* | 4484 | 11 910 | 54,388 |
|  |  | (606 - 23,889) | (1,215 - 256,980) | (208 - 140,087,698) |
|  | *Run 4* | 4411 | 11,026 | 44,596 |
|  |  | (532 - 29,134) | (1,208 - 153,462) | (85 - 132,647,784) |
| ***CNP*** | *Run 1* | 1,125 | 11,644 | 12,477 |
|  |  | (45 - 9,391) | (2,757 - 54,175) | (178 - 461,211) |
|  | *Run 2* | 1,021 | 10,637 | 10,879 |
|  |  | (50 - 9,080) | (2,553 - 44,668) | (237 - 360,081) |
|  | *Run 3* | 726 | 11,048 | 7,497 |
|  |  | (12 - 8,819) | (2,704 - 47,446) | (71 - 368,468) |
|  | *Run 4* | 566 | 10,615 | 5,288 |
|  |  | (8 - 7,518) | (2,540 - 44,056) | (59 - 219,685) |
| ***CLNP*** | *Run 1* | 1,225 | 11,416 | 7,874 |
|  |  | (17 - 16,885) | (2,532 - 51,523) | (64 - 587,895) |
|  | *Run 2* | 972 | 10,583 | 6,734 |
|  |  | (22 - 12,951) | (2,566 - 43,823) | (80 - 446,375) |
|  | *Run 3* | 647 | 10,950 | 4,759 |
|  |  | (8 - 11,051) | (2,719 - 45,394) | (42 - 219,281) |
|  | *Run 4* | 534 | 10,397 | 3,612 |
|  |  | (4 - 10,566) | (2,523 - 42,717) | (24 - 289,468) |
| ***DNP*** | *Run 1* | 7,401 | 9,543 | 13,110 |
|  |  | (70 - 367,113,200) | (390 - 2,669,931) | (4 - 627,769,194) |
|  | *Run 2* | 3,921 | 9,356 | 32,085 |
|  |  | (38 - 187,932) | (364 - 284,512) | (21 - 693,904,973) |
|  | *Run 3* | 3,642 | 10,651 | 24,992 |
|  |  | (13 - 1,236,517) | (348 - 4,369,181) | (8 - 757,355,877) |
|  | *Run 4* | 2,769 | 10,610 | 17,600 |
|  |  | (8 - 109,144) | (370 - 2,441,181) | (10 - 571,610,240) |
| ***BNP*** | *Run 1* | 24,643 | 8000 | 6,473 |
|  |  | (1,057 - 1,244,514,612) | (120 - 4,305,266) | (4 - 1,316,739,913) |
|  | *Run 2* | 7,925 | 7,320 | 47,435 |
|  |  | (295 - 856,446) | (104 - 316,082) | (7 - 1,201,434,227) |
|  | *Run 3* | 8,126 | 8,547 | 85,645 |
|  |  | (168 - 5,794,287) | (50 - 14,151,419) | (6 - 2,553,877,136) |
|  | *Run 4* | 6,716 | 8,555 | 75,875 |
|  |  | (43 - 1,085,425) | (55 - 13,696,197) | (5 - 2,552,701,303) |

Figure S5 Results for 10 bootstraps for each individual PSMC analyses

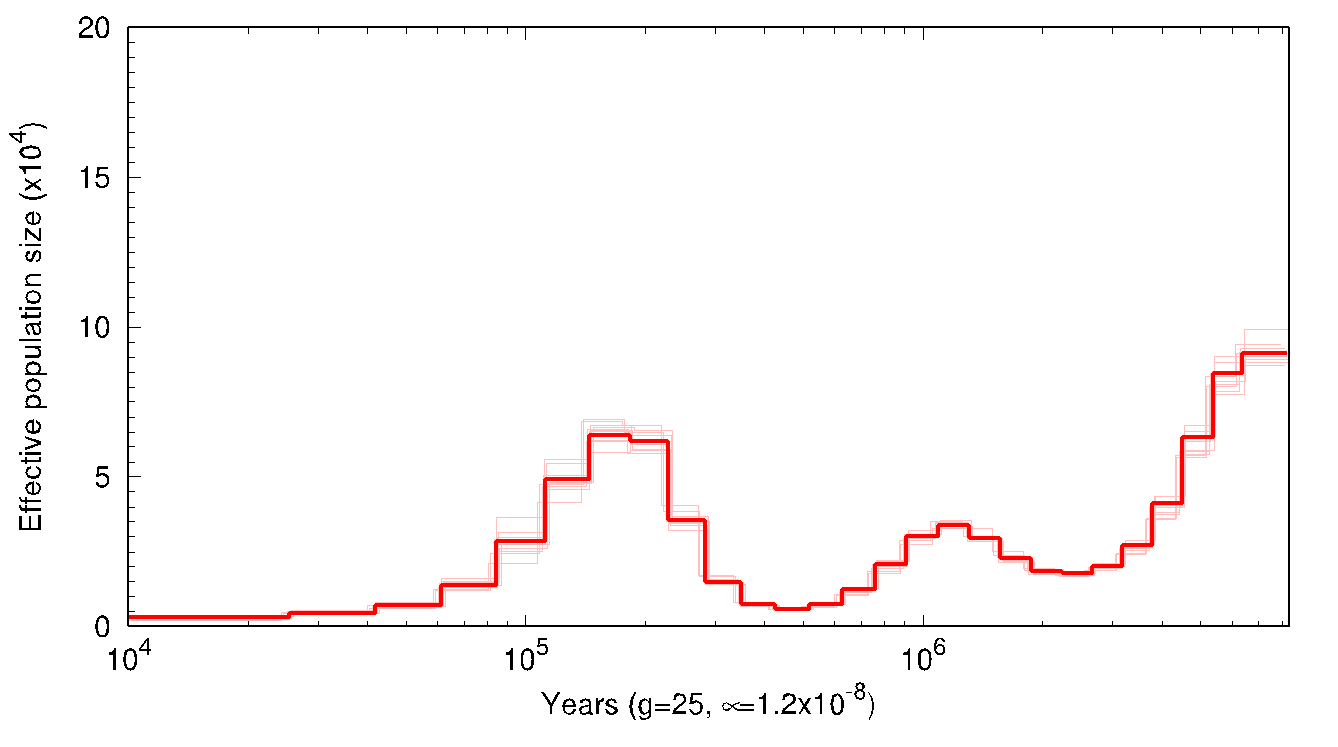

a) Bella

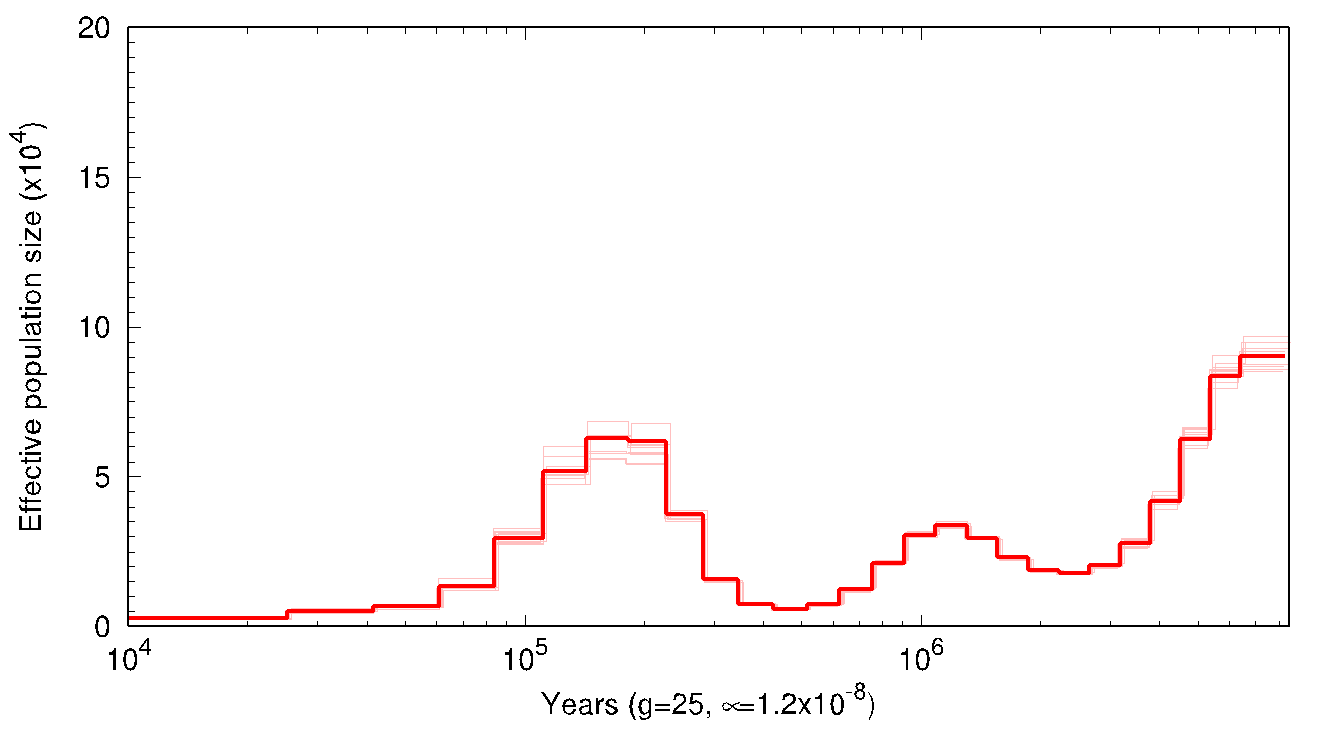

b) Bo

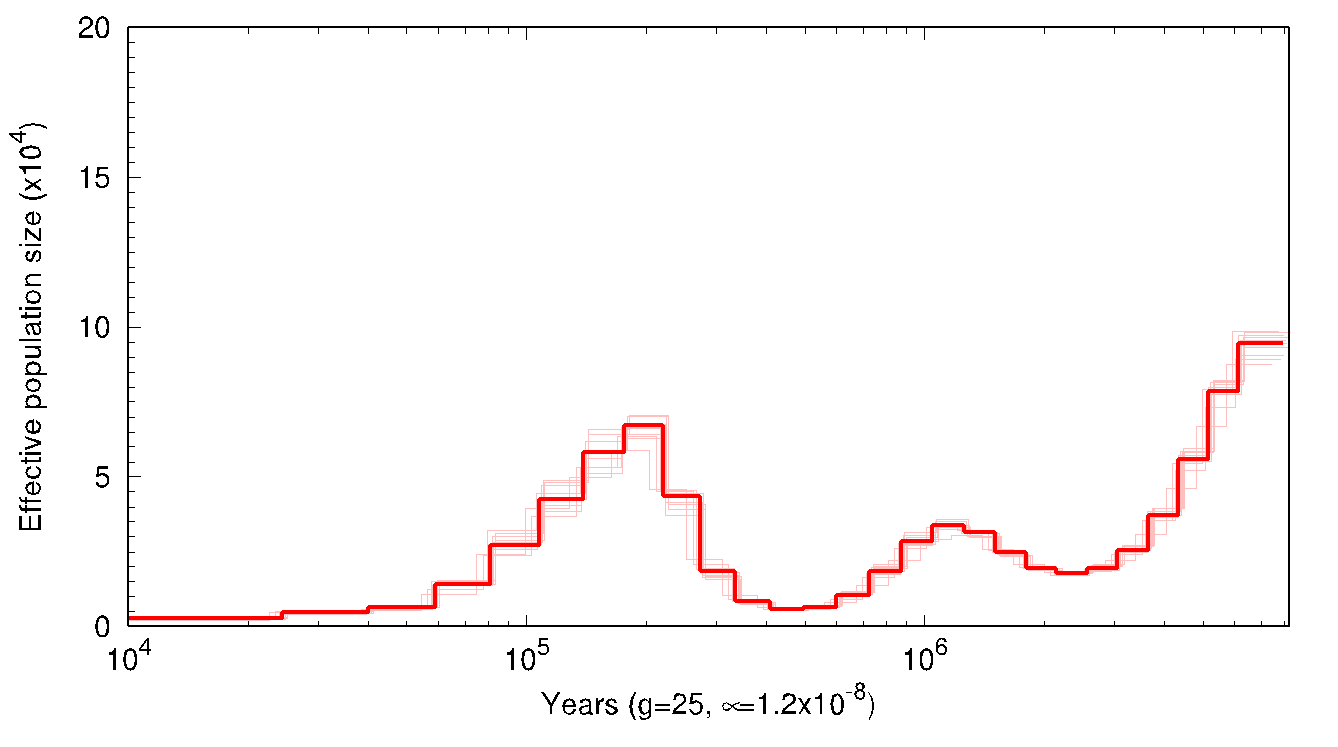

c) Emilia

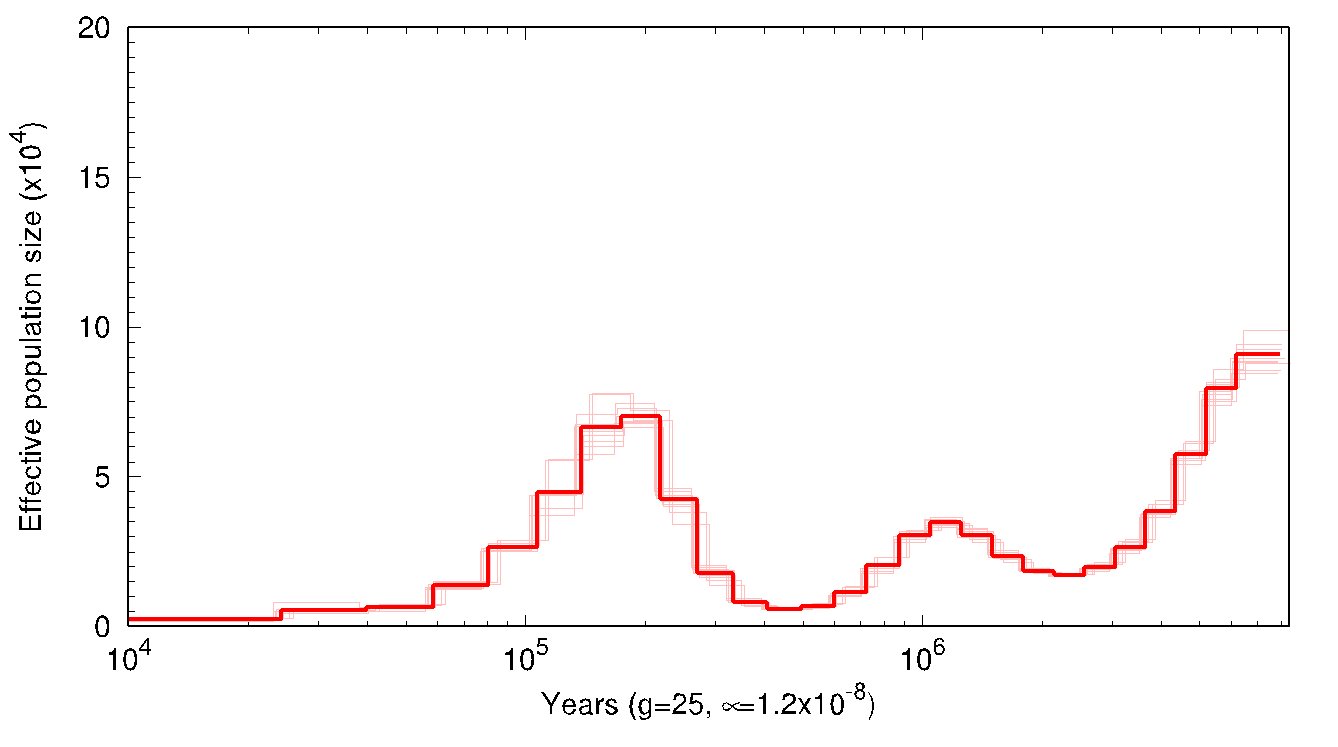

d) Simão

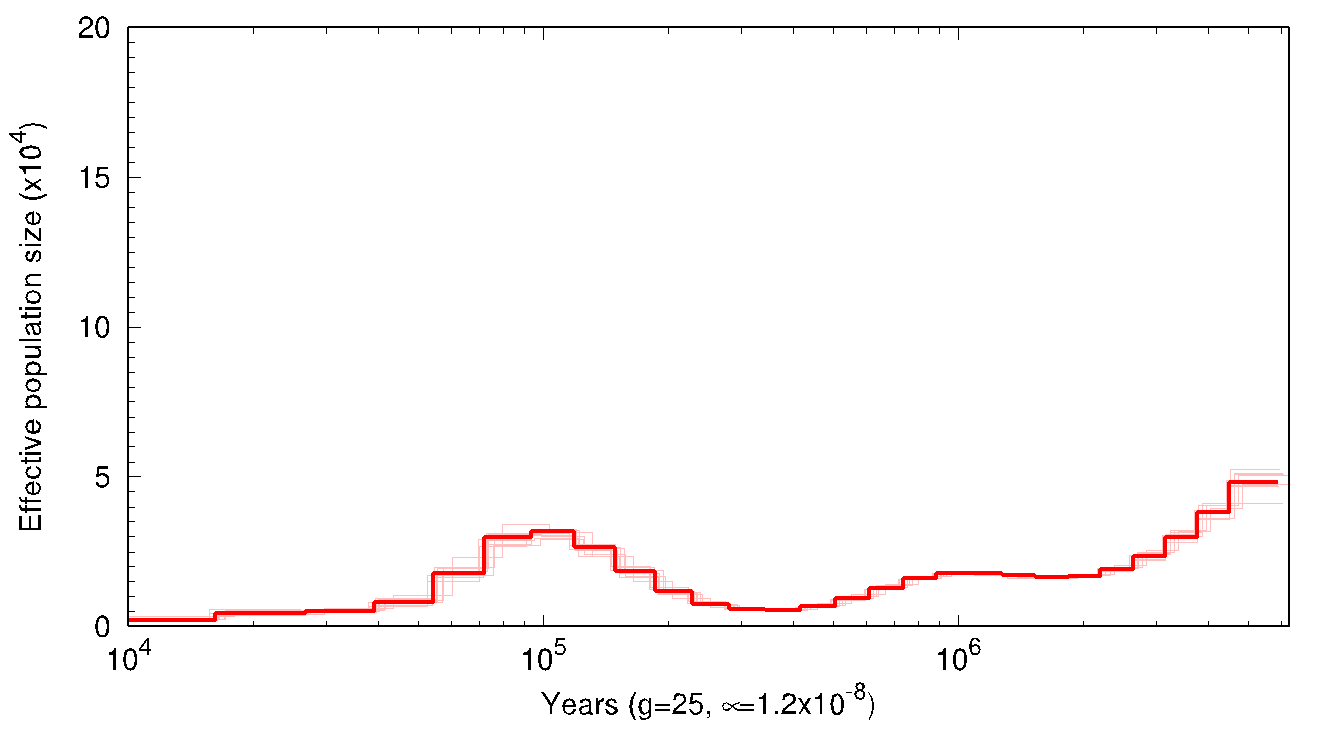

e) T-3_Chimp
